# Long range regulation of transcription scales with genomic distance in a gene specific manner

**DOI:** 10.1101/2024.07.19.604327

**Authors:** Christina L Jensen, Liang-Fu Chen, Tomek Swigut, Olivia J Crocker, David Yao, Mike C Bassik, James E Ferrell, Alistair N Boettiger, Joanna Wysocka

## Abstract

While critical for tuning the timing and level of transcription, enhancer communication with distal promoters is not well understood. Here we bypass the need for sequence-specific transcription factors and recruit activators directly using CARGO-VPR, an approach for targeting dCas9-VPR using a multiplexed array of RNA guides. We show that this approach achieves effective activator recruitment to arbitrary genomic sites, even those inaccessible by single dCas9. We utilize CARGO-VPR across the *Prdm8-Fgf5* locus in mESCs, where neither gene is expressed. We demonstrate that while activator recruitment to any tested region results in transcriptional induction of at least one gene, the expression level strongly depends on the genomic distance between the promoter and activator recruitment site. However, the expression-distance relationship for each gene scales distinctly in a manner not attributable to differences in 3D contact frequency, promoter DNA sequence or presence of the repressive chromatin marks at the locus.

## Introduction

In multicellular organisms, the spatiotemporal control of transcription is regulated by non-coding sequences scattered throughout the genome known as enhancers. Canonically defined as functioning independently of the genomic distance and orientation to their target promoter^1^, enhancers are bound by transcription factors (TFs), which in turn recruit coactivator complexes that mediate transcriptional activation through a variety of mechanisms^2–5^. Enhancers can be located at a vast range of genomic distances from 1kb to over a megabase^6–8^ from the gene they regulate. However, the mechanism by which active enhancers communicate with their target promoters across large genomic distances and through three-dimensional space remains elusive despite considerable interest in the field. While previous models of enhancer-promoter (E-P) communication evoked formation of stable chromatin loops, recent reports suggest that enhancer-promoter contacts are highly dynamic and relatively infrequent^9–11^. Furthermore, while some imaging studies have documented clear associations between E-P proximity and transcription^11–13^, others reported a lack of temporal correlation between the two^10,14^. A model reconciling these seemingly disparate observations has been proposed, whereby the relationship between E-P contact frequency and transcription is nonlinear. This nonlinearity arises from transient E-P interactions indirectly leading to promoter activation through a probabilistic, multi-step process that temporally decouples E-P contact and transcription^15,16^.

Regardless of the specific model, the emerging view is that, however transient or indirect, E-P contacts are required for enhancer-activated transcription. Such contacts can be facilitated by the process of loop extrusion by cohesin that results in genome folding into ‘topologically associating domains’ (TADs)^17,18^. TADs partition the chromosome into ∼100 kb to 2 Mb regions of self-interacting chromatin that are insulated from other regions by boundary elements, bound by a zinc finger protein CTCF^17,18^. Enhancers often regulate promoters that are located within the same TAD, although examples of the TAD boundary bypass by enhancers also exist ^19–21^. Contacts also arise intrinsically as a function of genomic distance owing to the polymeric nature of chromatin which specifies that contact frequencies between pairs of loci decays sharply with their genomic distance, with a power law scaling^22^. Given the dependence of transcription (directly or indirectly) on E-P contacts and the relationship between contacts and genomic distance, it follows that an enhancer’s effect on promoter transcription should be dependent on the E-P genomic distance. Some existing evidence supports this dependency. For example, genomic distance is a good predictor of which promoter will be activated by an enhancer ^23,24^. Moreover, inserting a strong enhancer at different positions relative to the promoter shows that enhancer action diminishes with genomic distance^15,25–27^. Together, these observations raise a question of how the relationship between E-P distance and transcription scales at different loci.

Enhancer activity is highly constrained by the dependence on recognition by transcription factors whose binding motifs are encoded in the enhancer sequence. However, it is unclear to what extent features beyond enhancer sequence – such as its position within the broader regulatory domain – enable enhancer competence. One way to decouple enhancer position in the genome from its sequence is through direct recruitment of coactivators in a sequence agnostic manner. Previous studies demonstrated that the requirement for native sequence-specific transcription factors (TFs) at enhancers can be bypassed by the synthetic recruitment of strong activation domains or specific coactivators to the enhancer DNA^14,28–30^. In particular, catalytically inactive Cas9 (dCas9) fused to activation domains such as VP64, VPR or to the coactivator p300 has been shown to achieve gene activation at long-ranges^24,30–33^. Indeed, VP64 and VPR robustly bind multiple coactivators such as Mediator and p300/CBP, providing a mechanistic explanation for their strong activation properties^34^. We reasoned that such synthetic activator recruitment via dCas9, when applied systematically across a regulatory domain, could provide a DNA sequence-agnostic approach to address two outstanding questions: (i) are all distal regions within a specific regulatory domain competent to activate gene expression upon activator recruitment?, and (ii) how does transcription from the promoter scale with the genomic distance to the distally-recruited activator?

To probe these questions, we use RNA-guided recruitment of dCas9-VPR to different regions of a regulatory domain containing *Prdm8* and *Fgf5* genes, both of which are transcriptionally inactive in mouse embryonic stem cells (mESCs). We first demonstrate that dCas9 targeting to promoter-distal regions using single guide RNAs often fails to result in detectable dCas9 binding, thereby confounding interpretation of negative results. We show that this limitation can be overcome using CARGO – a multiplexed array of gRNAs^35^ – to recruit dCas9-VPR (CARGO-VPR) efficiently to arbitrary genomic regions, including previously inaccessible chromatin sites. We then deploy CARGO-VPR to probe long-range activation of *Fgf5* and *Prdm8* promoters upon activator recruitment to many sites across the *Prdm8*-*Fgf5* locus. We show that although activator recruitment across the TAD results in measurable transcriptional induction of at least one gene, the level of activation is strongly dependent on the distance of the promoter from the activator recruitment site. Interestingly, however, this expression-distance relationship scales distinctly for the two genes within the TAD. We then show that the distinct behaviors of the two genes in response to distal activator cannot be explained by differences in contact frequency decay with genomic distance. Finally, neither swapping the two promoters nor disrupting repressive marks at the locus changes the expression-distance scaling relationship of the two genes, indicating that this relationship is not simply explainable by the inherent sequence properties of the promoters or the presence of repressive marks. Together these results demonstrate that arbitrary genomic sites within a regulatory domain can function like enhancers upon activator recruitment, but that the magnitude of transcriptional response is highly dependent on the genomic distance from the promoter. Moreover, our data suggests that different genes can integrate such activating signals distinctly, and that these differences are encrypted beyond promoter sequence.

## Results

### dCas9-VPR tiling screen at the *Prdm8-Fgf5* locus

We envisioned that using dCas9-VPR to recruit coactivators to any chosen genomic region in mESCs would enable us to distinguish between two possibilities: (i) any site to which coactivators are recruited can function as an enhancer, provided it makes sufficient contact with the promoter, or (ii) only certain regions within a regulatory domain are competent to activate distal promoters upon coactivator recruitment (**Figure 1A**). We chose the *Prdm8-Fgf5* locus for this work because the two genes are inactive in mESCs and fall within a compact (∼160 kb) TAD, flanked by two strong CTCF sites (**Figure 1B**)^36^. Moreover, *Fgf5* expression is induced during the transition from mESCs to the developmentally related cell state: epiblast-like cells (EpiLCs) while *Prdm8* is not expressed in either cell state. Thus, although inactive in mESCs, the *Fgf5* promoter should be competent for activation. Furthermore, the enhancers that drive *Fgf5* induction during this transition have been well characterized, including E1-E4 distal enhancers that become active in EpiLCs, and a proximal poised enhancer (PE) which is already hypersensitive in mESCs but lacks H3K27ac^37,38^ (highlighted in **Figure 1B**).

**Figure 1.**
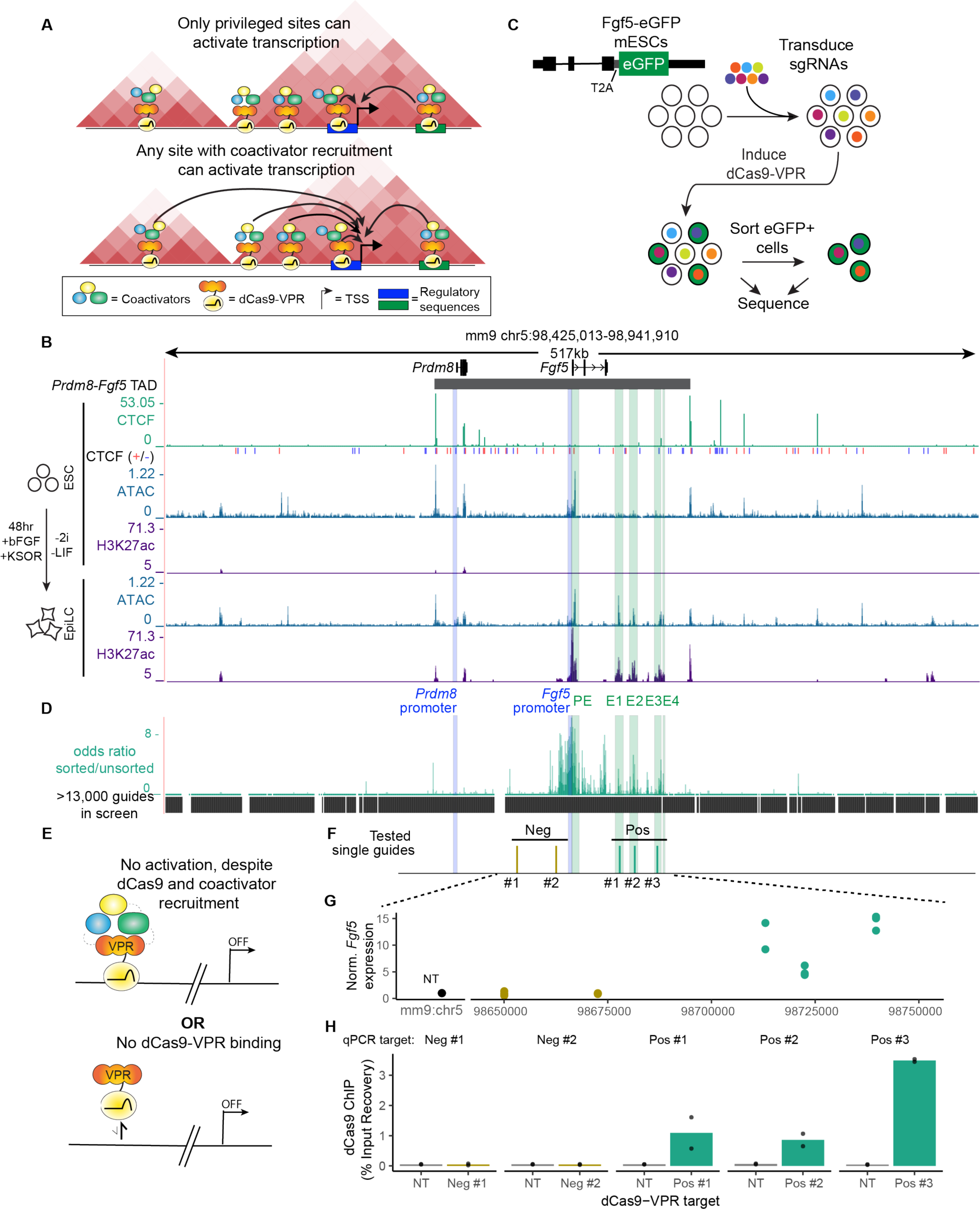
Lack of dCas9 binding confounds interpretation of CRISPRa screen. **A.** Schematic demonstrating two scenarios in which long-range activation is limited to privileged sites (upper) or is compatible with any site given sufficient cofactor recruitment (lower). **B.** Genome browser tracks of CTCF (ChIP-seq)^36^, CTCF motif orientations (+ strand, red,-strand, blue), H3K27ac (ChIP-seq)^37^ and ATAC-seq^43^ at the *Prdm8-Fgf5* locus in mESCs (top three tracks) and EpiLCs (bottom two tracks). TAD position is highlighted as grey bar at the top. **C.** Schematic of CRISPRa screen to identify regions capable of activating Fgf5-eGFP. **D.** Enrichment of guides in the eGFP+ population relative to the unsorted population (over 13,000 guides screened, lower black bars). **E.** Schematic of possible interpretations of negative CRISPRa results **F**. Positions of single guides tested for dCas9 binding. Gold bars indicate single guides unable to activate *Fgf5* in CRIPRa screen. Green bars are activating guides (enriched in the eGFP+ population). **G.** RT-qPCR of *Fgf5* expression in cells expressing dCas9-VPR and select single guides, as indicated in F. **H.** dCas9 ChIP-qPCR from cells expressing dCas9-VPR and select single guides (same guides as F and G).

To facilitate high-throughput readout of *Fgf5* expression by fluorescence activated cell sorting (FACS), we endogenously inserted a T2A-EGFP reporter at the end of the *Fgf5* gene in mESCs engineered to express doxycycline-inducible dCas9-VPR (**Figure 1C**). To guide dCas9-VPR recruitment, we designed a tiling library with >13,000 sgRNAs spanning 500kb centered at the *Prdm8-Fgf5* TAD (**Figure 1D**) and 1000 non-targeting control sgRNAs. We transduced the *Fgf5-T2A-EGFP*/dCas9-VPR mESCs with this library at a low multiplicity of infection to ensure that most cells received one or zero sgRNAs, induced dCas9-VPR expression, and sorted EGFP+ cells by FACS (**Figure 1C**). Subsequent sequencing and analysis of sgRNA enrichment in sorted vs unsorted populations uncovered sgRNAs which were able to activate *Fgf5* (**Figure 1D**). As expected, we observed that guides targeting dCas9-VPR at or near (<7kb) the *Fgf5* promoter were highly enriched in the EGFP+ population, in agreement with previous observations that CRISPRa is most efficient at or in proximity to promoters^39^. Interestingly, however, promoter-distal sgRNAs able to activate *Fgf5* preferentially target the *Fgf5* EpiLC enhancers. In contrast, the promoter-distal regions upstream of the *Fgf5* gene were conspicuously devoid of activating sgRNAs, as were regions flanking the *Prdm8-Fgf5* TAD (**Figure 1D**).

### Lack of dCas9 binding confounds interpretation of the CRISPRa screen

These observations suggest that enhancers that are inactive in mESCs but become accessible in a subsequent developmental state are privileged sites for long-range *Fgf5* activation, whereas many other sites within the *Prdm8-Fgf5* locus are not permissive. Similar observations were recently made in CRISPRa/i screens in hESCs, in which activating/repressing distal sgRNAs were limited to so-called ‘pre-established transcriptionally competent chromatin regions’. Many of these regions lacked hypersensitivity or active/poised enhancer marks but showed low levels of binding by TFs, histone deacetylases, and DNA demethylases^40^. However, an important factor confounding interpretation of our and other CRISPRa/i screens is that dCas9 binding is not measured. Thus, an alternative possibility is that the lack of activity for a specific region is reflective of the absence of dCas9 (and its fused effector domain) recruitment to the site, rather than incompetency of the targeted region for long range activation *per se* (**Figure 1E**). To test this, we first selected several guides that were either present in the eGFP enriched population (**Figure 1F**, green) or absent from it (**Figure 1F**, gold), generated stable lines expressing these guides and dox-inducible dCas9-VPR and measured expression of *Fgf5* by RT-qPCR. Consistent with the screen results, guides enriched in the eGFP population, and not those without significant enrichment, activated *Fgf5* expression (**Figure 1G**). We then performed ChIP-qPCR for dCas9 in these lines and observed dCas9 binding only at sites that were able to activate *Fgf5* (**Figure 1H**). Thus, single RNA guides are, in many cases, incapable of efficient recruitment of dCas9 to promoter-distal genomic regions, likely due to the fact that nucleosomes can impede dCas9 access to DNA^41^. However, it has also been shown that sporadic unwinding or ‘breathing’ of nucleosomal DNA facilitates Cas9 activity^42^. While the *Fgf5* EpiLC enhancers are not open in mESCs (as measured by ATAC-seq^43^), some of the TFs known to bind them in EpiLCs (e.g. Oct4 and Sox2) are also expressed in mESCs. Therefore, these TFs might sporadically engage *Fgf5* EpiLC enhancers, in turn facilitating nucleosome breathing, guide pairing with the underlying DNA and dCas9-VPR binding. Regardless of the specific mechanism at play, our observations indicate that negative results in CRISPRa screens cannot be interpreted without measurements of dCas9 binding, especially at closed chromatin sites.

### Targeting dCas9-VPR with multiple guides enables recruitment to previously inaccessible sites

To assess the long-range enhancer competence of arbitrary genomic regions we had to overcome the inability of dCas9 targeted by single guide RNAs to bind closed chromatin regions. We hypothesized that stable dCas9-VPR binding could be achieved by targeting dCas9 with multiple closely spaced guides (**Figure 2A**). We previously developed novel molecular assembly strategy, termed chimeric array of gRNA oligos (CARGO), that achieves highly multiplexed gRNA delivery into single cells^35^. To test if using CARGO arrays would increase binding of dCas9-VPR at regions inaccessible to single guides, we designed two sets of guides spanning two distinct ∼500bp regions at the *Prdm8-Fgf5* locus: (i) 12 guides 14kb upstream of the *Fgf5* promoter (14kb USF), and (ii) 11 guides 35 kb downstream of the *Fgf5* promoter (35kb DSF). We then assembled the first six, last six, or all guides targeting each region into CARGO arrays (**Figure 2B**, **Figure S2A**) and measured binding of dCas9 by ChIP-qPCR upon stable array integration into mESCs expressing inducible dCas9-VPR. We first examined dCas9 binding at the 14kb USF site using three different PCR amplicons, corresponding to the third, fifth and tenth guide target sites within the region (labeled A, B and C, respectively, in **Figure 2B**). We compared CARGO-targeted binding with that achieved with single guides and observed a robust increase in dCas9-VPR occupancy when recruited by CARGO arrays versus single guides (**Figure 2C**). As expected, dCas9-VPR binding at the A site was higher with the CARGO 1-6 array, targeting the left portion of the region, whereas dCas9-VPR binding at the C site was higher with the CARGO 7-12 array, targeting the right portion (**Figure 2C**). Interestingly, increasing the number of guides to 12 by combining the two 6-mers (CARGO 1-12) did not further increase dCas9 binding (**Figure 2C**). The dCas9-VPR occupancy was accompanied by an increase in expression of *Fgf5*, signifying that even though the interrogated regions were not detected as hits in our original screen, effective recruitment of activators enables these regions to induce *Fgf5* transcription from a distance (**Figure 2D**). Highly concordant results were observed with CARGO arrays vs single guides at the 35kb DSF site (**Figure S2B,C**). In summary, CARGO-VPR allows for effective dCas9-VPR recruitment to non-coding regions inaccessible with single guides, facilitating quantitative interrogation of long-range gene regulation upon activator recruitment to arbitrary genomic sites.

**Figure 2.**
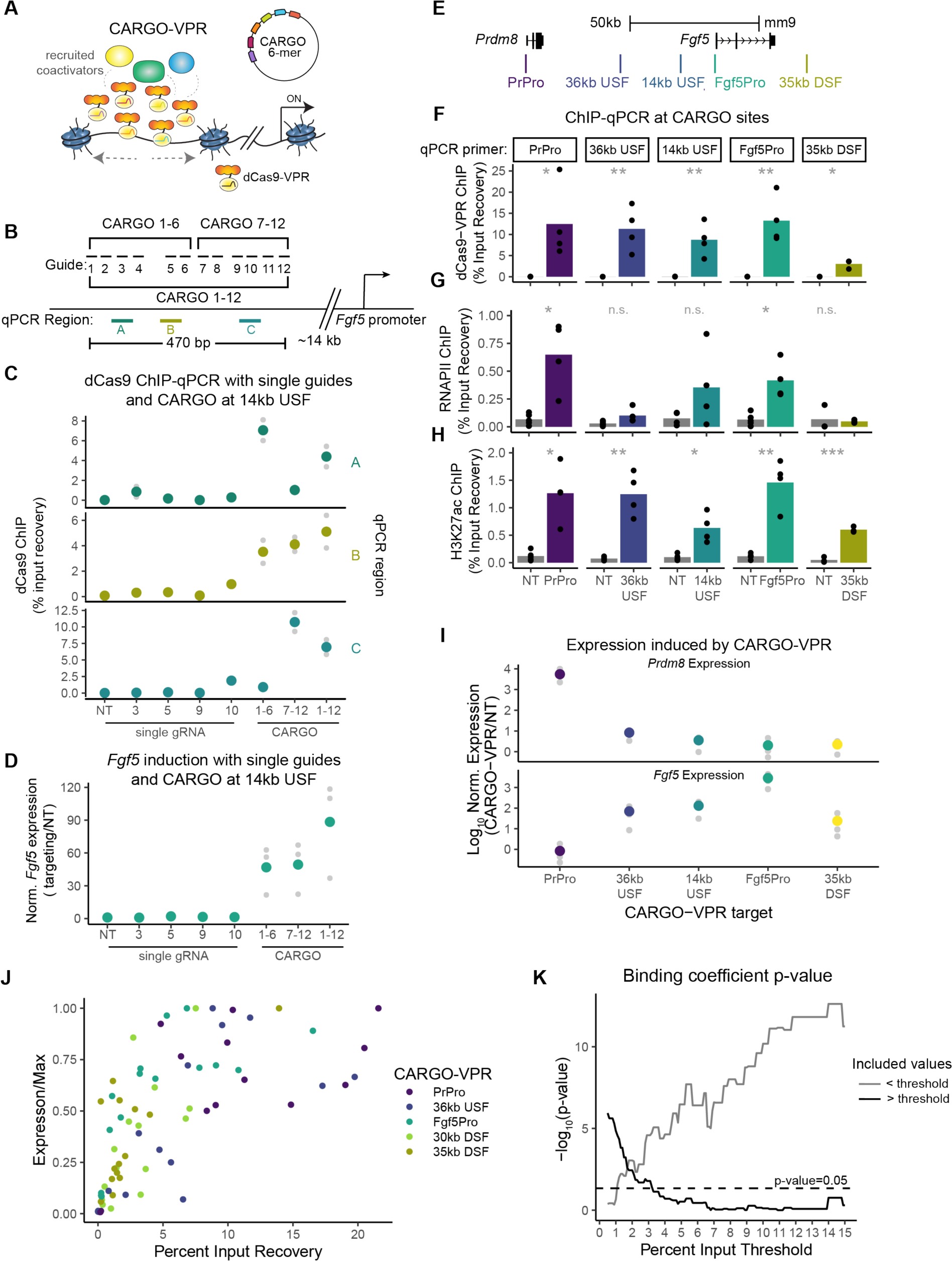
CARGO-VPR enables dCas9 recruitment to previously inaccessible sites. **A.** Schematic of CARGO-VPR strategy, utilizing a 6-mer gRNA arrays (CARGO arrays) for dCas9-VPR targeting to genomic regions of interest **B.** Positions of twelve individual guides, two 6-mer arrays, and one 12-mer array designed to target a region 14kb upstream of the *Fgf5* promoter (14 kb USF). Locations of PCR amplicons used in dCas9 ChIP-qPCR are shown as colored lines labeled A, B, and C. **C.** dCas9 ChIP-qPCR percent input recovery at sites labeled in (B) in cells expressing dCas9-VPR and single guides (non-targeting (NT), 3,5, or 10), 6-mer arrays (1-6 or 7-12), or a 12-mer array (1-12). Colored dots are the mean of two biological replicates and grey dots are the individual replicates. **D.** RT-qPCR of *Fgf5* expression in cells expressing dCas9-VPR and single guides (NT, 3,5, or 10), 6-mer arrays (1-6 or 7-12), or a 12-mer array (1-12). Colored dots are the mean of three biological replicates and grey dots are the individual replicates. **E.** Positions of 6-mer CARGO arrays at the *Prdm8*-*Fgf5* locus. **F.** dCas9, **G.** RNAPII, and **H.** H3K27ac ChIP-qPCR in cells expressing CARGO arrays shown in (E) or a NT guide. Individual panels correspond to a PCR amplicon overlapping indicated array. Bar height is the mean of n=3-5 replicates with each replicate shown as black dots (*=p<0.05,**=p<0.01, ***p<0.001, Welch’s two sample t-test). **I.** RT-qPCR of *Prdm8* (upper) and *Fgf5* (lower) in cells expressing indicated CARGO-VPR target. Expression shown as log_10_ ratio of normalized expression in the CARGO-VPR line over normalized expression in the NT control. Colored points are the mean of 4 replicates. Grey points are individual expression values. **J.** Expression normalized by max observed for each CARGO-VPR target plotted against percent input recovery in cells expressing dCas9-VPR and 1, 3, 4, 5, or 6 guides. **K.** negative log p-values for the binding coefficient in a linear regression model of expression as a function binding (percent input recovery) for models including binding data at different, indicated percent input recovery thresholds. See also Figure S2

### CARGO-VPR promotes H3K27ac enrichment at targeted regions

We next examined if CARGO-VPR results in gain of features indicative of active regulatory elements, such as RNAPII binding and H3K27ac. We assembled 6-mer CARGO arrays targeting regions corresponding to the promoters of *Prdm8* (PrPro) and *Fgf5* (Fgf5Pro) and three distal non-coding regions at the locus, located between the two genes (36kb USF and 14kb USF) or downstream from *Fgf5* (35kb DSF) (**Figure 2E**). We confirmed dCas9-VPR binding by ChIP-qPCR (**Figure 2F**). We analyzed RNAPII binding and H3K27ac at the targeted regions following dCas9-VPR induction, comparing ChIP-qPCR signals from cells expressing targeting or control non-targeting CARGO arrays. We observed elevated RNAPII occupancy at the promoters, but not at the distal sites (**Figure 2G**). H3K27ac levels were significantly enriched at all tested regions in cells expressing targeting arrays, consistent with gain of regulatory capacity (**Figure 2H**). Next, we analyzed ability of each CARGO array to activate *Prdm8* and *Fgf5* expression, as measured by RT-qPCR. We observed that targeting distinct regions at the locus resulted in considerably different transcriptional outcomes: promoter targeting arrays resulted in strong activation of their cognate gene but not of the gene expressed from the other promoter (**Figure 2I**). Furthermore, distal regions showed decreased ability to activate transcription (**Figure 2I**).

### Expression is only linearly dependent on activator binding when binding is low

Because we observed an increase in gene activation when we increased the number guides recruiting dCas9-VPR from one to six, we aimed to explore the relationship between dCas9-VPR occupancy levels and transcriptional activation in a context that is decoupled from distance effects. To this end, for three distal sites and two promoter sites, we generated a series of arrays in which we titrated the number of guides used to recruit dCas9-VPR (from 6 down to 5,4,3 or 1) and examined both dCas9 binding and gene expression (**Figure S2D**). In this way we were able to vary binding without changing the distance between promoter and activator recruitment site. We observed no or weak activation of expression with single guides at all tested sites. Increasing the number of guides resulted in increased activation of the target gene up to a saturation point. Expression induced by dCas9-VPR at either promoter reached a maximal level with only 3 to 4 guides, whereas the distal sites required more guides to approach maximal expression (**Figure S2E**). We observed that dCas9 occupancy generally increased with increased number of guides with little to no binding with single guides and detectable binding beginning with three guides used to recruit dCas9-VPR (**Figure S2F**).

Next, we plotted expression against dCas9-VPR percent input recovery in each of the paired samples. Expression increased sharply at low percent input recovery values, then typically plateaued, indicating that expression is only linearly dependent on binding at low binding levels (**Figue S2G**). To analyze the relationship between binding and expression across all sites in a combined manner, we normalized expression to the observed maximal expression value for each CARGO-VPR target site (**Figure 2J**). To determine the level of binding at which expression is no longer significantly predicted by binding, we examined linear regression models of the form Expression/Max ∼ B_1_binding + B_0_ run on a subset of the data either above or below the indicated percent binding threshold (**Figure 2K**). We observe that binding is predictive of expression when percent input recovery values below 3.5% are included. However, the relationship between expression and binding is no longer significant above this threshold.

### Expression from *Fgf5* and *Prdm8* promoters is highly dependent on distance from the activator recruitment site, but with distinct distance decay profiles

To further investigate the relationship between position within the TAD, E-P distance and gene expression, we designed and assembled 38 6-mer CARGO arrays spanning the entire *Prdm8-Fgf5* TAD, including several arrays outside of the TAD boundary (**Figure 3A**). We avoided targeting arrays within the body of each gene, to preclude interference of dCas9 with RNAPII passage. We stably integrated each array into clonal tetO dCas9-VPR mESC cell lines and induced dCas9-VPR expression for 3 days, at which point cells were harvested for further analyses (**Figure 3B**). In each of the 38 CARGO-VPR cell populations we measured dCas9-VPR binding at the respective site by ChIP-qPCR using two independent primer sets and confirmed strong occupancy at most sites, albeit at variable levels (**Figure 3C and S3A**). We then measured *Prdm8* and *Fgf5* expression by RT-qPCR and observed that all CARGO arrays activated at least one promoter above the non-targeting control, consistent with the notion that any region across the locus is competent for long-range gene activation upon coactivator recruitment (**Figure 3D**). However, we also observed a striking decay in levels of expression as we targeted sites further from the promoter (**Figure 3D and E**). We observe no detectable differences between the upstream and downstream expression decay with distance at each promoter even when considering arrays that lie outside the TAD upstream of *Prdm8* (**Figure 3D and E**). To ensure that these results are robust to clonal variation or small changes in dCas9-VPR expression levels, we reproduced our observations using two different clonal tetO dCas9-VPR mESC lines for derivation of the CARGO-VPR cell populations, with concordant results (**Figure S3B and C**).

**Figure 3.**
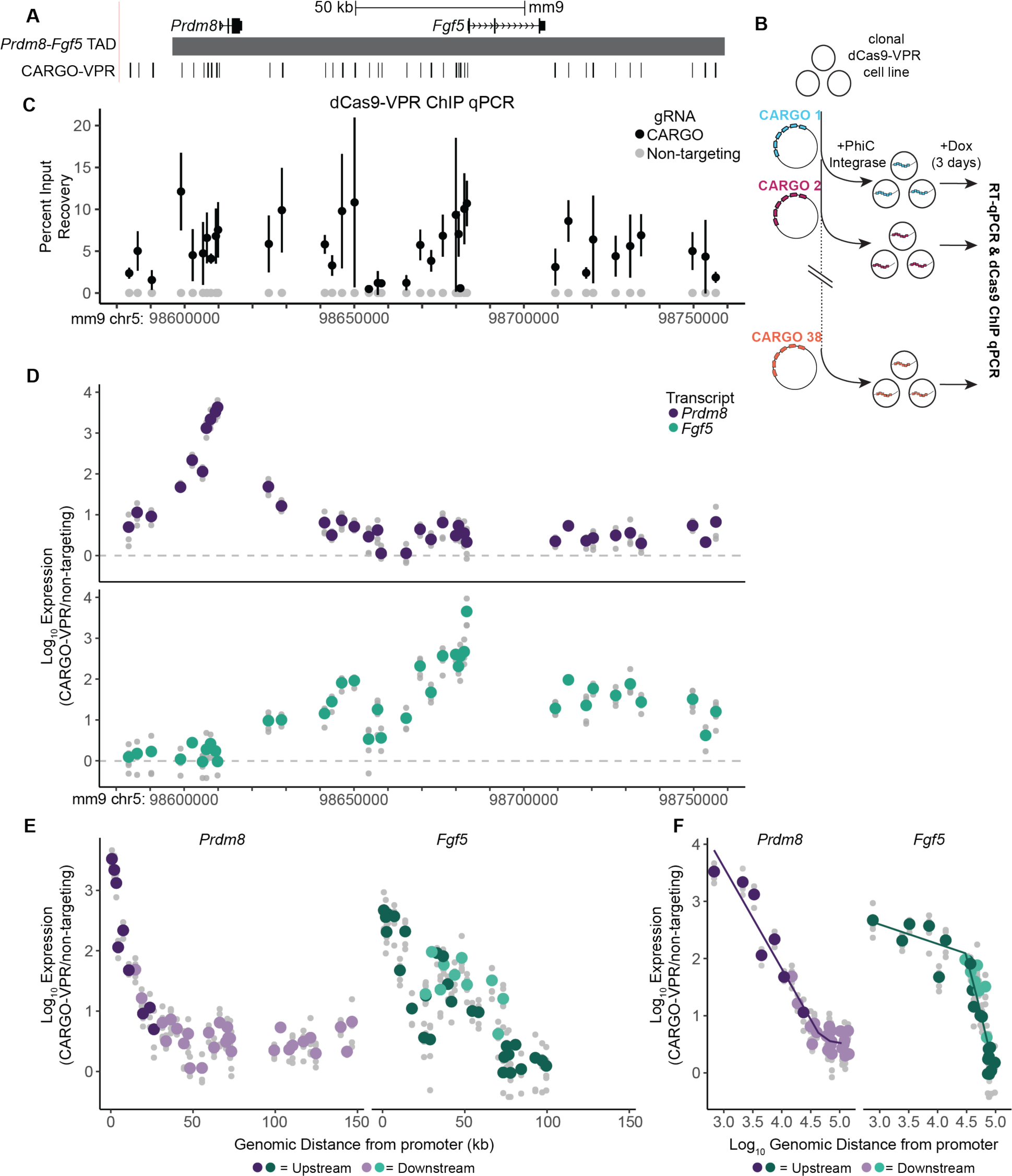
Gene-specific decay profiles of expression with genomic distance. **A.** Positions of 38 CARGO-VPR arrays targeting the *Prdm8-Fgf5* locus. **B.** Schematic of CARGO-VPR line generation. After dCas9-VPR clonal lines are isolated, each CARGO array is stably integrated using PhiC integrase generating 38 cell populations. Each population is treated with doxycycline for three days prior to downstream analyses such as RT-qPCR and ChIP-qPCR. **C**. dCas9 ChIP-qPCR in all 38 CARGO-VPR cell populations. Black points are the mean of n= 3-4 biological replicates. Error bars are standard deviation. Grey points indicate the same qPCR amplicon assayed from a cell population expressing a non-targeting guide (n=2). **D.** RT-qPCR of *Prdm8* (upper, purple) and *Fgf5* (lower, green) from cell populations expressing dCas9-VPR and a CARGO array targeted to each of the positions shown on x-axis. Colored dots are the mean of at least four replicates. Grey points are individual expression values. Expression is plotted as expression normalized to a housekeeping gene, *Rpl13a*, over normalized expression from a population expressing a non-targeting guide. **E.** RT-qPCR of *Prdm8* (Left) and *Fgf5* (Right) plotted against absolute distance of array from the target promoter. Data points from arrays located upstream or downstream from the promoter are shown as triangles and dots, respectively. Colored dots/triangles are the mean of at least four replicates. Grey points are individual expression values. Expression is plotted as expression normalized to housekeeping gene, *Rpl13a*, over normalized expression from a population expression a non-targeting guide. **F.** Log_10_ RT-qPCR expression in CARGO-VPR lines normalized to expression in non-targeting line plotted against log_10_ distance of CARGO-VPR recruitment site and promoter. Arrays that have an average percent input recovery less than 3.5% were excluded from analysis. Spline regression fits with slopes of -1.77 and -0.18 for *Prdm8* and -0.34 and -4.30 for *Fgf5* (adjusted R^2^ 0.94, 0.87,respectively). See also Figure S3

Interestingly, the expression levels of both genes have distinct distance decay profiles, with *Fgf5* expression showing a shallower decay with distance compared to that of *Prdm8* (**Figure 3D and E**). To explore the nature of this difference, we aimed to estimate the scaling relationship between expression of each gene and distance to the activator. To minimize the effects of variable dCas9-VPR binding on this analysis, we removed arrays that were measured to have a mean percent input recovery of <3.5%. This is consistent with our observation that 3.5% represents the threshold above which greater binding does not contribute significantly to higher expression values (**Figure 2K**). If a power law scaling relationship exists between distance and expression for either of the genes, we expect to observe a linear relationship on a log-log plot with a slope corresponding to the scaling factor. Plotting the log-scaled expression as a function of log-scaled genomic distance (**Figure 3F**), we observe that *Prdm8* expression has a constant scaling factor with respect to distance up to a distance of about 50kb at which point we cannot detect changes in *Prdm8* expression above background levels (**Figure 3F**, slope = -1.77). However, *Fgf5* expression does not have a single scaling parameter with respect to distance, but rather has two regimes of distance dependence. When activators are targeted within ∼15-30kb of the promoter, *Fgf5* expression is buffered against increasing distance. Targeting beyond this buffered zone leads to *Fgf5* expression falling sharply with distance (**Figure 3F**, buffered slope = -0.34, responsive slope = -4.30). Together, our data demonstrate that while expression of both genes at the locus is highly dependent on the distance from the activator recruitment site, the distance decay expression profiles are distinct for each gene. Our results further suggest the key feature of the distinction between the two genes is that *Fgf5* expression is buffered against increasing activator distance up ∼15-30kb whereas *Prdm8* expression has little or no such buffering.

### 3D contact frequencies do not explain distinct expression profiles

We next asked whether distinct expression-distance relationships could be explained by differences in 3D contacts made by the two promoters. For example, if *Fgf5* promoter contacts the surrounding chromatin more frequently than does the *Prdm8* promoter, this might account for the observed differences in expression scaling in response to activator recruitment. To test whether distinct features in the 3D structure of the *Prdm8-Fgf5* locus could be explanatory of differences in expression profiles, we turned to Optical Reconstruction of Chromatin Architecture (ORCA), a multiplexed sequential fluorescence in situ hybridization (FISH) method to image the *Prdm8-Fgf5* TAD using a series of probes tiling the locus at the 5 kb intervals ^9^ (**Figure 4A, B**). We imaged thousands of alleles using 38 custom probes targeted to the *Prdm8-Fgf5* locus. Prior to data analysis, we excluded probes that were unable to be detected in 50% or more alleles (6 of 38 probes) as well as alleles with fewer than 20 probes detected. Missing data was then imputed as an average from neighboring probes. This method enabled us to recover data for all points except at probes 32 and 33 (white stripe, **Figure S4A**).

**Figure 4.**
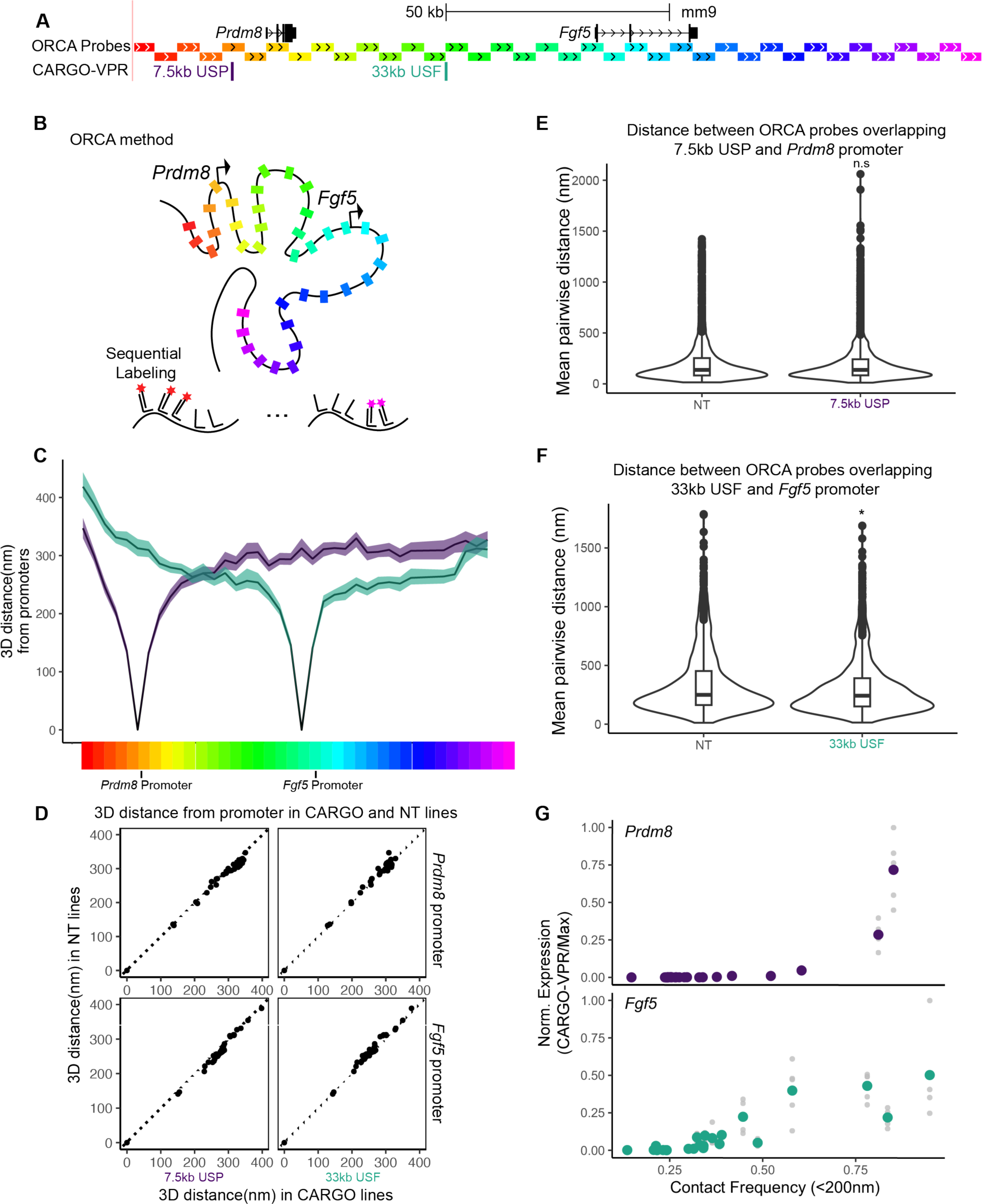
3D contact frequencies do not explain distinct expression profiles. **A.** Map of 38 5kb ORCA probes in relation to *Prdm8* and *Fgf5* and selected CARGO arrays that activate *Prdm8* (Purple, 7.5kb USP) or *Fgf5* (Green, 33kb USF). **B.** Schematic of ORCA strategy **C.** Pairwise median distance plots from the probes overlapping the *Prdm8* promoter or the *Fgf5* promoter (black lines) and all other probes. Location of individual probes is shown as rainbow-colored blocks and corresponds to placement in (A). Purple and green shading represent 95% confidence interval of the median. **D.** Median pairwise distance from probe overlapping *Prdm8* promoter (upper) or *Fgf5* promoter (lower) and other probes in cells expressing a non-targeting guide compared with equivalent median pairwise distances in cells expressing CARGO arrays in (D). Dotted lines are y=x. **E.** Distribution of pairwise distances between probes overlapping the *Prdm8* promoter and the 7.5kb USP CARGO-VPR site in dCas9-VPR cells expressing a non-targeting guide (NT) or 7.5kb USP CARGO (7.5kb USP). **F.** Distribution of pairwise distances between probes overlapping the *Fgf5* promoter and the 33kb USF CARGO-VPR site in dCas9-VPR cells expressing a non-targeting guide (NT) or 33kb USP CARGO (33b USF) **G.** Expression normalized to max observed of *Prdm8* (upper) and *Fgf5* (lower) induced by CARGO-VPR arrays with strong binding (>3.5%) (y-axis) by contact frequency (x-axis) between the promoter and the CARGO target site, as extrapolated from ORCA measurements. Colored dots are the mean of at least four replicates. Grey points are individual expression values. See also Figure S4

We first examined the population-average topology by looking at median pairwise inter-probe distances from hundreds of individual chromatin fiber traces across the entire TAD (**Figure S4A**). We observed constant scaling between median 3D distance and genomic distance across the TAD, with lack of strong intra-TAD features (**Figure S4A**). We note that this is in contrast to previously published Micro-C data from mESCs grown in serum, which show a condensed region around the *Prdm8* gene body and contacts between the two promoters (**Figure S4B**)^44^. Differences in media conditions-namely, 2i+Lif used here versus serum-have been shown to alter H3K27me3 deposition in cells and thus may potentially account for the observed differences in locus topology^45^. In agreement, analysis of the published H3K27me3 ChIP-seq datasets from two distinct mESC lines grown in 2i+Lif or serum, revealed an enrichment of H3K27me3 across the *Prdm8* gene body and both promoters in cells grown in serum as compared to those grown in 2i+Lif (**Figure S4C**)^45^.

We next investigated the median 3D distance measured between the probes that bind each promoter and all other probes in cells expressing dCas9-VPR and a non-targeting guide. At both promoters, we observed similar, symmetric distribution of 3D distance changes in relation to genomic distance (**Figure 4C**). Congruent results were obtained when contact frequencies of the two promoters were compared instead, calculated as the frequency of each probe coming into functional contact (<200nm) with each promoter **(Figure S4D**). We reasoned that it may be possible that activation of *Prdm8* or *Fgf5* might alter the conformation of the region. Thus, we performed ORCA in mESCs expressing CARGO-VPR that activates either *Prdm8* or *Fgf5* (**Figure 4A**). Comparing pairwise distances between promoters and all probes in activated versus non-targeting lines, we observed a high degree of correlation between samples (**Figure 4D**). We next examined the pairwise 3D distances between the promoters and the CARGO-VPR recruitment sites in activated and non-targeting lines (**Fig 4E and F**). We observe no significant difference in the distance distributions between the CARGO-VPR recruitment site (7.5kb USP) and the *Prdm8* promoter in activating versus non-targeting lines (**Fig 4E**). We do observe a significant, but small difference in the distance distributions of the CARGO-VPR recruitment site (33kb USF) and the *Fgf5* promoter in the activating versus non-targeting lines (Mann-Whitney U test, p = 0.015, NT median distance: 250 nm, activating CARGO median distance: 243 nm) (**Fig 4F**). We reasoned that if this this small change in median distance was due to the emergence of a fraction of alleles with the CARGO-VPR recruitment site being more frequently in proximity to the promoter, we would observe a more bimodal distribution of these distances. However, using Hartigans’ dip test for unimodality, we see no evidence of multimodality (p = 0.94-0.99) in any of the distributions plotted here, even upon activation. Together, these results indicate that activation of either gene by CARGO does not substantially alter overall locus topology or 3D distance between the promoter and the activator recruitment site.

To further explore the relationship between 3D contacts and expression, we extrapolated contact frequency at each CARGO-VPR site determined to have strong binding (>3.5%) from the contact frequency measured by ORCA (see Methods for details). We observed that, as reported previously^15,16^, the relationship is non-linear, namely, that small changes in contact frequency result in large changes in expression (**Figure 4G**, plotted on semi-log scale in **S4E**). However, the two promoters at the locus show distinct responses, with *Fgf5* expression being induced and reaching maximal expression at lower contact frequencies than *Prdm8* expression (**Figure 4G**), consistent with *Fgf5* having a buffered regime in regions with frequent contacts or close genomic distance. Altogether, we do not observe differences in 3D contacts that would explain the distinct expression distance decay profiles of the two promoters, but rather the response to contacts frequency is gene-specific.

### Repressive chromatin marks do not explain distinct expression profiles

We considered that presence of repressive marks at the locus may alter expression responses of the two genes. Although, as shown in **Figure S4C**, there is substantially less H3K27me3 at the *Prdm8-Fgf5* locus in 2i+Lif compared to serum conditions, this mark is overall more enriched over *Prdm8* gene in either condition. Moreover, there is an 8kb block of repressive chromatin marked by H3K9me3 at a full-length long terminal repeat (LTR) element located downstream from the *Prdm8* gene (**Figure S5A**)^46^. To test if the presence of the heterochromatic LTR element affects scaling relationships at the locus, we deleted the LTR in the dCas9-VPR cells, integrated CARGO arrays and measured expression. We observed almost no change in the *Prdm8* or *Fgf5* expression profiles between Wild type and the LTR deletion lines (**Fig S5B and C**). To test if loss of repressive H3K27me3 mark changes the *Prdm8* promoter response, we inhibited activity of the Polycomb Repressive Complex 2 (PRC2) with an allosteric inhibitor EED226. Treatment with EED226 was sufficient to diminish H3K27me3 levels at the *Prdm8* promoter as well at downstream sites with lower H3K27me3 enrichment (**Figure S5D**). However, when we examined *Prdm8* responses to long-range activation by CARGO-VPR following treatment with this drug, we observed remarkably similar profiles to that of cells treated with vehicle control (**Figure S5E**). Together, these results suggest that the repressive chromatin marks at the locus do not account for the scaling differences we observe between the two genes.

### Promoter sequence is insufficient to alter scaling of expression responses

It has been proposed that the non-linearity between E-P contact frequency and expression is a consequence of a multi-step process determined by the promoter state and its activation/deactivation kinetics^15^. We therefore hypothesized that distinct expression scaling of the two promoters at the *Prdm8-Fgf5* locus might be a consequence of differences in their cis-regulatory features. To examine if differences in expression responses of the two promoters are determined by their intrinsic sequence features, we swapped the *Prdm8* and *Fgf5* promoters (corresponding to the region beginning 350bp or 400bp upstream of each TSS and ending 100bp or 50bp downstream) by inserting the *Fgf5* promoter in place of that of *Prdm8* and inserting the *Prdm8* promoter in place of the *Fgf5* promoter (**Figure 5A and S5F)**. We then generated a new set of cell lines expressing each of the CARGO arrays in this promoter swap genetic background. Following induction of dCas9-VPR, we performed RT-qPCR from *Prdm8* and *Fgf5* RNA driven by the exogenous swapped promoter and compared it to the RNA levels measured in WT dCas9-VPR lines expressing identical CARGO arrays. We observed a striking correspondence in the expression profiles between the two genotypes, with the only outliers being associated with the promoter-targeting CARGOs (as expected, since the promoters are now swapped) (**Figure 5B and C**). Together, these observations indicate that differences in the chromatin state and sequence of the two promoters at the *Prdm8-Fgf5* locus are insufficient to explain their distinct expression scaling in response to the same distal activator.

**Figure 5.**
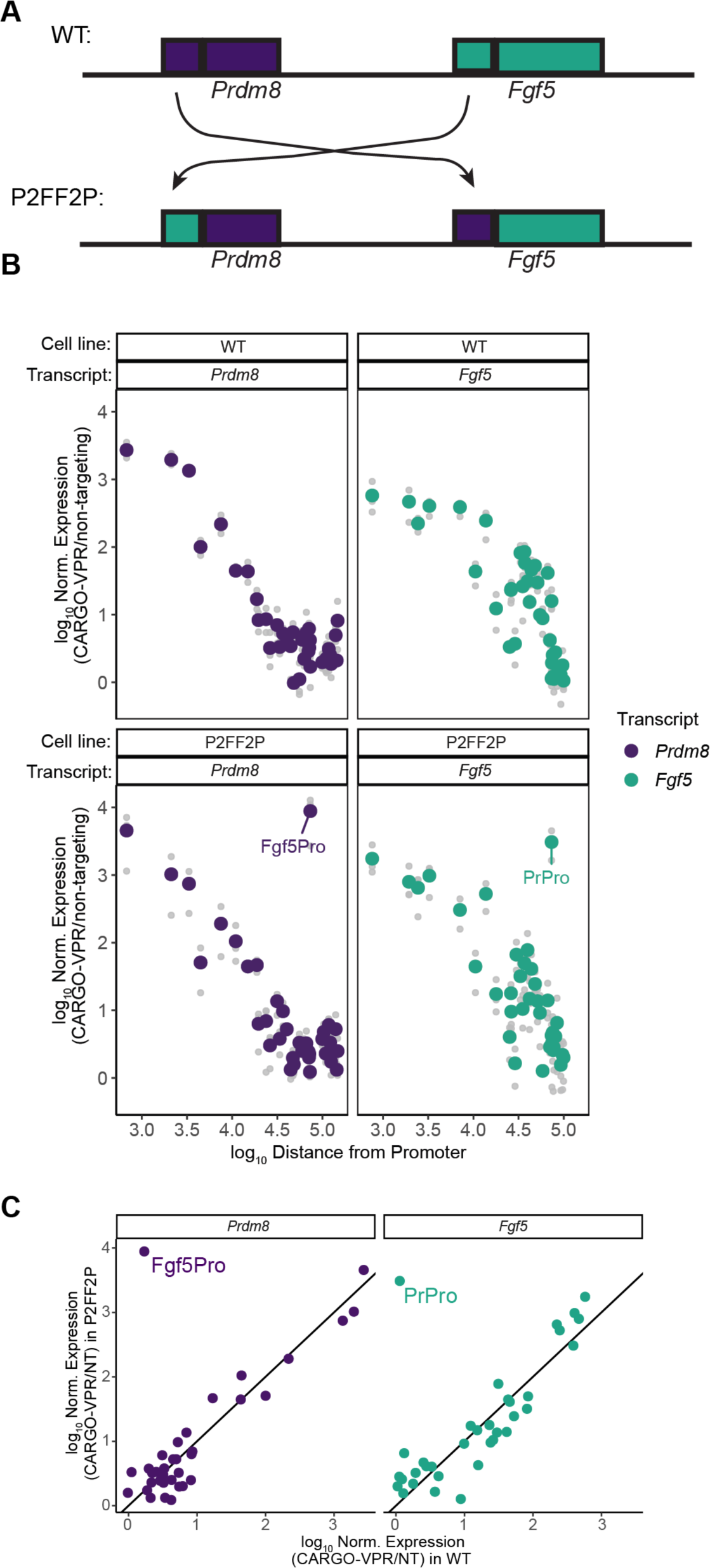
Promoter swap does not alter scaling of expression responses. **A.** Schematic of the *Prdm8* and *Fgf5* promoter swap. 450bp sequence surrounding each TSS was swapped in P2FF2P mESCs (see Methods for details). **B.** RT-qPCR expression of *Fgf5* and *Prdm8* in WT and promoter swap (P2FF2P) lines expressing dCas9-VPR and CARGO arrays. Colored points are the mean of three replicates. Grey points are individual expression values. **C.** Comparison of *Fgf5* and *Prdm8* expression levels in promoter swap (P2FF2P) vs WT mESCs upon CARGO VPR. line is y=x. Points are mean of three biological replicates. See also Figure S5

## Discussion

Our work describes CARGO-VPR as an approach to interrogate long-range regulatory capacity of arbitrary genomic regions. We pinpoint paucity of dCas9 binding as a critical limitation in CRISPRa screens of promoter-distal, closed chromatin sites, and show that this issue can be circumvented by targeting such regions with multiplexed gRNA arrays. The assay described here is complementary to the previously reported strategy of locally mobilizing an enhancer via transposon system to test effects of numerous positions relative to a promoter^15^. CARGO-VPR has an added advantage of being able to reliably detect potential negative events – namely those resulting in detectable activator recruitment to the site, but no effects on expression. Our approach could be expanded in future studies to test whether any site of interest is competent for activation and – by recruiting distinct dCas9 fusion proteins – to examine how specific coactivators, corepressors or their enzymatic subunits impart gene regulation at long-ranges.

Using CARGO-VPR we assessed regulatory capacity of diverse regions across the *Prdm8-Fgf5* locus and studied quantitative relationship between genomic distance and gene expression. Interestingly, all CARGO-VPR targeted regions within the examined regulatory domain showed ability to activate at least one promoter, suggesting that any non-coding genomic region might be in principle compatible with enhancer function, provided it assembles coactivators and has sufficient contact with a transcriptionally-permissive promoter. Furthermore, our work indicates that long-range activation decays with the E-P genomic distance, as has also been demonstrated by Zuin et al.^15^. A parsimonious explanation for these observations is that enhancer-promoter contacts are required for activation, and since contacts between pairs of genomic loci on the chromatin polymer decay with distance, so does the activation of transcription. These observations raise a question of how certain enhancers are capable of activating transcription at ultra-long ranges of a megabase of DNA or more^6–8^. One potential explanation is that other processes, such as loop extrusion by cohesin or presence of tethering elements near enhancer and promoter, are necessary to counteract the effects of genomic distance and increase contact frequency between such ultra-long range E-P pairs ^26,47,48^. In agreement with the ultra-long range enhancers being distinctly reliant on loop extrusion, cohesin is required for activation of *Shh* by enhancers located many hundreds of kilobases away, but dispensable for those located within 100 kb from the target gene^29^.

Although enhancer action decreases with distance, the relationship between contact frequency and transcription is non-linear^15^. While our results are consistent with this non-linearity, they also emphasize that the relationship between contact frequency and transcription is gene specific, even within the same TAD. We expected that the promoter would be the main integration site of the activator inputs, and thus that the differences in the promoter sequence would drive observed differences in the expression-distance scaling relationship at *Prdm8* and *Fgf5*. Surprisingly, however, when we swapped the two promoters, we observed that the distinct expression decay profiles are not dependent on the promoter itself, despite selecting a generous promoter region that includes proximal regulatory sequences. Similarly, depleting H3K27me3 from the *Prdm8* promoter did not alter the expression-distance scaling. Given that the contact frequency also decays equivalently with distance at the two promoters, our observations suggest that other characteristics of each gene beyond the promoter contribute to the distinct distance decay profiles. At present, we do not know what these characteristics are, as depleting repressive chromatin marks had no effect on the scaling relationship. There are also no active mESC enhancers at the locus and no strong CTCF sites, beyond those at the TAD boundaries. We do note the presence of weaker CTCF sites downstream from the *Prdm8* gene, but both our ORCA data and analysis of the expression induced by the equidistant CARGO arrays on both sides of the *Prdm8* promoter are inconsistent with those sites serving as a boundary. The only distinguishing feature at the locus is the presence of an open but inactive poised enhancer downstream of the first exon of *Fgf5*. Previous work showed that during EpiLC transition, the poised enhancer is activated and amplifies the output of other, more distal *Fgf5* enhancers^49^. Thus, one possibility remains that even in mESCs, the poised enhancer amplifies the effect of CARGO-VPR, resulting in a dampened decay of expression with distance.

A key characteristic of the observed gene-specific differences is that *Fgf5* expression is buffered against the distance-dependent decay when targeting activators to sites up to 15-30kb away, whereas *Prdm8* has no (or very little) buffering. This buffering zone could result from the saturation of contacts, such that once contact frequencies above a certain threshold are reached, the maximal expression is achieved. Nonetheless, such saturation likely does not simply reflect exhaustion of the promoter’s capacity for activation, because targeting CARGO-VPR directly at the promoter results in higher expression than that observed in the buffered zone for distal activation. Rather, the buffering might represent saturation of a step preceding transcription initiation, such as E-P communication. While the biochemical mechanism of E-P communication, that is, the identity of the message relayed from the enhancer to the target promoter to activate transcription, remains unclear, our data are consistent with a model in which each gene has a distinct radius of ‘functional’ contact. Sites that are close in genomic distance to the promoter will reside within this functional radius more frequently than sites that are further away. For *Fgf5*, we observe a larger radius of functional contact such that targeting within 15-30kb in genomic distance achieves maximal communication whereas *Prdm8* does not have such a buffered zone and targeting just 5kb away from the promoter is marked by a distance dependent decay in expression. Several biochemical mechanisms could be consistent with this model such as activator condensates or activator hubs localized to regulatory elements. Regardless of the particular underlying mechanism, our results show that the scaling of expression with enhancer distance is gene specific and that genomic context beyond the promoter must contribute to this specificity.

## Limitations of the Study

While we demonstrate that recruitment of CARGO-VPR to promoter-distal sites results in local H3K27ac enrichment and achieves long-range activation of gene expression, we acknowledge that CARGO-VPR recruitment to arbitrary sites may not be fully equivalent to activation mediated by endogenous enhancers. Further, it is conceivable that long residency times of dCas9 may alter the dynamics of activation compared to that mediated by the native transcription factors.

## Acknowledgements

We thank members of the Wysocka lab for critical reading of the manuscript. We thank Seungsoo Kim for the modified version of the pk335-BFP plasmid and Kristel Dorighi for sharing EpiLC ATAC-seq data. This work was supported by HHMI (to J.W.), R35 GM131757 (to J.W.), U01 DK127419 (to J.W. and A.N.B.), Lorry Lokey endowed professorship (to J.W.), Beckman Young Investigator Award (to A.N.B.), Packard Fellowship for Science and Engineering (to A.N.B.), National Science Foundation Graduate Research Fellowship (to C.L.J.).

## Author Contributions

Conceptualization, C.L.J, T.S., and J.W; methodology C.L.J., L.F.C., O.J.C., D.Y.; analysis C.L.J., L.F.C., T.S.; visualization C.L.J; writing-original draft C.L.J and J.W., writing-review-all authors; funding C.L.J, J.W., A.N.B; Mentorship J.W., T.S., A.N.B, J.E.F.; Supervision J.W.

## Resource availability

### Lead contact

Further information and requests for resources and reagents should be directed to and will be fulfilled by lead contact, Joanna Wysocka

### Materials availability

Plasmids generated in this study will be deposited to Addgene.

### Data and code availability

- All data reported in this paper will be shared by the lead contact upon request.
- All original code is available in this paper’s supplemental information.
- Any additional information required to reanalyze the data reported here is available from the lead contact upon request.

## Experimental model and subject details

### Culture of mouse embryonic stem cells

R1 mESCs were cultured as previously described^50^. Briefly, male R1 cells were grown in monolayer on tissue culture plates pretreated with 7.5 ug/ml poly-L-ornithine (Sigma) in PBS and 5 ug/ml laminin (Gibco) in PBS consecutively for at least 1 hour at 37°C. Cells were maintained in serum-free 2i + LIF media (500mL DMEM/F-12 (Gibco), 1x Gem21 Neuroplex w/o Vitamin A (Gemini Bio), 1x N2 Neuroplex (Gemini Bio), 2.5g BSA (Gemini Bio), 1xMEM NEAA (Thermo), 1xSodium pyruvate (Thermo), and 1xAnti-Anti (Sigma) containing MEK inhibitor PD0325907 (0.8 uM, Selleck), GSK3β inhibitor CHIR99021 (3.3 μM, Selleck), and leukemia inhibitory factor (LIF).) Cells were passaged when 80-90% confluent using accutase (Sigma) to release cells from the plate. Cells were re-plated on PLO/Laminin plates at 1:6-1:12 dilutions.

### Method details

#### Endogenous tagging of *Fgf5*

mESCs were transfected using Lipofectamine 2000 (Thermo) per manufacturers protocol with px458-mCherry carrying Cas9 and a single sgRNA targeting the C-terminus of *Fgf5* and a donor template containing LoxP-UBC promoter-Neomycin resistance-LoxP-t2a-EGFP flanked by 1kb homology arms that were amplified from genomic DNA. Cells were treated with 200ug/ml G418 (Invivogen) then sorted as single cells by FACS onto fibronectin (Thermo) treated 96-well plates. Homozygous clones were identified by PCR. A homozygous clone was then transfected as above with a plasmid with Cre-recombinase to remove the loxP-UBC-Neomycin selection cassette leaving only a LoxP scar. Cells were single cell sorted by FACS as above and homozygous clones were identified using PCR and sanger sequencing (Elim).

### Generating doxycycline-inducible dCas9-VPR clonal lines

PiggyBac dCas9-VPR (addgene #191269) was digested with BsrGI-HF (NEB) to remove a fragment containing tagBFP and a portion of dCas9. To regenerate the complete dCas9 sequence, primers # and # were used to amplify the dCas9 fragment and Gibson assembly was used to insert this sequence back into the BsrGI-HF digested backbone resulting in PiggyBac dCas9-VPR without tagBFP.

mESCs were transfected using lipofectamine 2000 (Thermo) per manufactures protocol with PiggyBac dCas9-VPR (noBFP) and PiggyBac transposase. Cells were treated with 3ug/mL blasticidin (Invivogen) to select transformants then were plated on PLO/Laminin treated 10cm plates at 1000 cells/plate and colonies were isolated.

### sgRNA library construction

The gRNA library was PCR amplified from a pooled-oligo chip (CustomArray, Agilent) using Herculase II Fusion Polymerase (Agilent, 600675), then cloned into a lentiviral gRNA expression vector pMCB320 (https://www.addgene.org/89359/) using the restriction enzymes BstXI (NEB, R0113S) and BlpI (NEB, R0585S) and T4 DNA Ligase (NEB, M0202T). Ligation products were purified using QIAGEN MinElute columns (QIAGEN, 28004), then electroporated into Endura Electrocompetent Cells (Biosearch Technologies, 60242-1) using a Bio-Rad Gene Pulser Xcell system (Bio-Rad, 1652662) set at 1.8 kV, 600 ohms, and 10 uF in a 0.1 cm cuvette (Bio-Rad, 1652089). Cultures were spread onto LB-Agar (Fisher Scientific, DF0445-17-4) plates with Carbenicillin (Fisher Scientific, BP2648250) to maintain a minimum of 100X bacterial coverage per library element, then grown at 30C overnight before harvesting and extracting the final plasmid library using the QIAGEN HiSpeed Plasmid Maxi Kit (QIAGEN, 12643). Guide sequences are provide in **Table S2**.

### CRISPRa tiling screen

Lentivirus harboring tiling sgRNA library including 1000 non-targeting sgRNAs was produced as previously described in 293FTs. Mouse embryonic stem cells harboring the endogenous EGFP tag in place of the Fgf5 stop codon were transduced with lentivirus on 15 cm plates with 2 ug/ml polybrene. 20hr after transduction, viral media was replaced with fresh 2i+LIF media containing 2 ug/ul doxycycline and 1ug/ul puromycin. After 4-5 days of selection, cells were released with Trypsin, washed with PBS and resuspended in cold FACS buffer (2% FBS, 2 mM EDTA, 1% Sodium Azide in PBS). Cells were sorted based on positive EGFP signal on a FACS Aria II flow cytometer (BD) in the Stanford Stem Cell FACS facility. Four batches of sorted and unsorted cells were collected, pelleted and snap frozen. Genomic DNA from sorted and unsorted cells was harvested using the Gentra Puregene kit (Qiagen). Libraries for each sample were generated as previously described. Briefly, total genomic DNA was amplified by PCR with Huclease II polymerase using oMCB_1562/oMCB_1563 guide-specific primers (**Table S1**). Like samples were pooled and used as template for a second PCR in which Illumina adaptors (oMCB_1439) and barcodes were added to the samples. Samples were pooled for sequencing on the Miseq (Illumina).

### Guide enrichment analysis

Guide enrichment was quantified using quasibinomial regression in R. Briefly, presence in the sorted or unsorted populations were modeled with guide position (ID) and batch as predictors. The odds ratio between the test guides and negative guides was extracted from this model as the exponentiated coefficient. Multiple comparison correction was applied with an FDR of 0.1. Guides and results are provided in **Table S2**.

### Cloning CARGO arrays

CARGO arrays were generated as described previously^35^. Guide RNA sequences and custom oligos for CARGO synthesis are provided in **Table S3**. CARGO constant region was extracted after digestion from pGEMT-hU6-SL using BsmBI (NEB) at 55°C for 4 hours followed by gel purification (Machery-Nagel). CARGO plasmid backbone was generated by sequential digest of pK335 BFP plasmid DNA with XbaI at 37°C overnight followed by PciI at 37°C for 4hours followed by gel purification (Machery-Nagel). Custom oligos (10um each) were annealed and phosphorylated in 1x T4 ligase buffer (NEB) with 2.5 units T4 Polynucleotide Kinase (NEB, M0201) at 37°C for 30 minutes, 98°C for 5 minutes then temperature was decreased at a rate of -0.1 °C/s until a final temperature of 16 °C.

Each anneal oligo was combined with the digested CARGO constant region to generate minicircles. Briefly, 96ng CARGO constant region and 8.32uL of 50x diluted annealed oligos were ligated in 1x T4 ligase buffer (NEB) with 8 Weiss units T4 DNA ligase (Thermo) at room temperature for two hours. Minicircles were then treated with Plasmid-safe exonuclease to remove unligated fragments according to the manufacturer’s protocol. Minicircles were then combined and cleaned using Zymo DNA Clean & Concentrator-5 kit and eluted in 8uL Molecular Bio grade water.

CARGO array was then generated from the minicircles and the CARGO plasmid backbone. Briefly, minicircles and backbone were combined at a 3:1 ratio in 1x Tango buffer (Thermo) containing 0.10uL of 0.1M DTT, 0.40uL of 25mM ATP, T7 DNA ligase (NEB), and 10units BpiI. Samples were then incubated in a thermocycler for 50 cycles of 37°C for 5 minutes/20°C for 5 minutes. Assembly reaction was then treated with Plasmid-safe exonuclease per the manufacturer’s protocol. 1uL of this reaction was used to transform 25uL NEB Stable competent *E.coli*.

### Generating single guide and CARGO populations

Clonal R1 mESCs expressing doxycycline-inducible dCas9-VPR and blasticidin-resistence gene were transfected using lipofectamine 2000 (Thermo) per manufacturer’s protocol with 1:1 (w/w) PhiC Integrase plasmid and either pk335 with single guide or CARGO array. 24 hours post transfection, cells were treated with 200ug/mL G418 (Invivogen) and 3ug/ml blasticidin (Invivogen). Populations were passaged at least twice in G418+blasticidin media before being used for downstream experiments.

### ChIP

For all ChIP experiments, cells were treated with 2ug/ul doxycycline (Sigma) for three days to induce dCas9-VPR expression. At least two biological replicates from distinct cell passages were collected. For H3K27me3 ChIPs following EED226 (Selleck) treatment, treatment plates were treated with 10uM EED226 and 2ug/ul doxycycline and control plates were treated with 0.1% DMSO (vehicle control) and 2ug/ul doxycycline for three days.

After three days of treatment, cells were washed twice with PBS then fixed using freshly prepared 1% methanol-free formaldehyde in PBS solution. Cells were incubated on an orbital shaker for 5 minutes at room temperature (RT), then glycine was added to a final concentration of 0.125M and incubation continued for 5 min. The quenched formaldehyde solution was removed, and cross-linked cells were scraped from plate in ice cold PBS with 0.001% triton-X100. Scraped cells were transferred to a suitable tube and pelleted at 1350g for 5 minutes at 4°C, washed with ice cold PBS, and pelleted as before. All liquid was removed then pellets were snap-frozen in liquid nitrogen and stored at -80°C until further processing.

Cell pellets were thawed on ice for 30 minutes then resuspended in 5ml ice cold LB1 (50 mM HEPES-KOH pH 7.5, 140 mM NaCl, 1 mM EDTA, 10% glycerol, 0.5% NP-40, 0.25% Triton X-100, with 1X Roche cOmplete Protease Inhibitor Cocktai1 (PIC) and 1mM PMSF), rotated vertically at 4°C for 10 minutes, then pelleted at 1350g for 5 minutes at 4°C. Lysates were resuspended in 5mL ice cold LB2 (10mM Tris-HCl, pH 8.0, 200mM NaCl, 1mM EDTA, 0.5mM EGTA with 1X PIC and 1mM PMSF), rotated vertically at 4°C for 10 minutes, then pelleted at 1350g for 5 minutes at 4°C. Lysates were then resuspended in 300ul ice cold LB3 (10mM Tris-HCl, pH 8.0, 100mM NaCl, 1mM EDTA, 0.5 EGTA, 0.1% Na-Deoxycholate, 0.5% N-laurylsarcosine with 1x PIC and 1mM PMSF) and transferred to Diagenode TPX tubes and incubated on ice for 10 minutes. Lysates were sonicated for 12 cycles of 30s on/ 30s off on high power using Bioruptor Plus (Diagenode). Sonicated lysate was diluted to 1mL in LB3 and spun at max speed (>12000g) for 10 minutes at 4C. Supernatant was transferred to a fresh DNA LoBind tube and triton-X100 was added to a final of 1%.

To quantify DNA in each sample and enable equal DNA loading for immunoprecipitation, 10uL of each sample was taken for DNA extraction while remainder of samples were kept on ice. Briefly, to each 10uL sample, 90uL elution buffer (1% SDS, 0.1M NaHCO_3_), 2ul 5M NaCl, and 1ul RNaseA (0.2mg/ml final) was added. Samples were incubated at 65°C for 1hour with shaking. 1ul ProteinaseK (0.2mg/ml final) was added and samples were incubated at 65°C for an additional 1hour with shaking. DNA was then extracted using Zymo ChIP Clean & Concentrator-5 kit according to the manufacture’s protocol. DNA was quantified using a Nanodrop.

Remaining chromatin was then normalized and 50ul and 10ul aliquots for input samples were reserved and stored at -20°C. For each immunoprecipitation, 5uL antibody was added to each 1mL soluble chromatin sample, then samples were rotated vertically overnight (12-16hr) at 4°C.

On the next day, 100uL Protein G Dynabeads (Thermo) per ChIP sample were washed 3 times with ice cold blocking buffer (0.5% BSA(w/v) in PBS) then ChIPs were added to beads and incubated with vertical rotation for 4-6 hours at 4°C. ChIPs were then washed 5 times with RIPA buffer(50mM HEPES-KOH, pH 7.5, 500mM LiCl, 1mM EDTA, 1% NP-40, 0.7% Na-Deoxycholate), once with TE + 50mM NaCL (50mM Tris-HCl, ph 8.0, 10mM EDTA, 50mM NaCl), then eluted from the bead in 200uL elution buffer at 65°C for 30min. eluate was then transferred to a new DNA LoBind tube. To reverse cross-links and degrade RNA, 8uL 5mM NaCl and 2uL RNaseA (0.2mg/ml final) was added to eluate and to thawed input samples diluted to 200uL in elution buffer. All samples and inputs were incubated overnight (12-16hr) at 65°C with shaking. ProteinaseK (0.2mg/ml final) was added and samples and inputs were incubated for 2 hours at 55°C with shaking. DNA was extracted using Zyno Clean& Concentrator-5 kit. DNA was then stored at -20°C or used directly for ChIP-qPCR.

DNA samples were diluted by a factor of 8. Master mixes containing 5uL/reaction Sensifast SYBR No-Rox 2x mix (Meridian Bioline, 98020), 0.25ul 10mM primer mix, and 0.75uL water were prepared and kept on ice. 6uL of appropriate qPCR master mix was added to each well of a 384 well plate followed by 4uL of diluted cDNA. All samples were run in technical triplicate. qPCR was then run on a LightCycler 480 (Roche).

### RT-qPCR

For all RT-qPCR experiments, cells were grown in six-well plates, treated with 2ug/ul doxycycline for three days to induce dCas9-VPR expression. At least three biological replicates from distinct cell passages were collected. For experiments involving EED226 (Selleck) treatment, cells were plated in duplicate. The treatment well was treated with 10uM EED226 and 2ug/ul doxycycline and control wells were treated with 0.1% DMSO (vehicle control) and 2ug/ul doxycycline for three days prior to harvest.

Cells washed twice with PBS at RT then 750uL Trizol reagent (Invitrogen) was added to each well. After incubation on orbital shaker for 5 minutes at RT, samples were transferred to RNase free tubes and stored at -80°C.

To extract RNA, samples in Trizol were thawed on ice, then total RNA was extracted using Zymo Direct-zol kit according to manufacturer’s protocol. cDNA was prepared from 2ug RNA using High Capacity cDNA Reverse Transcription Kit (Thermo) according to manufacturer’s protocol with the exception that OligodT 20mer (IDT) was used in place of random primers and 1uL RNseOUT (Invitrogen) was added.

cDNA samples were diluted by a factor of 15 prior to RT-qPCR. Master mixes containing 5uL/reaction Sensifast SYBR No-Rox 2x mix (Meridian Bioline, 98020), 0.25ul 10mM primer mix, and 0.75uL water were prepared and kept on ice. 6uL of appropriate qPCR master mix was added to each well of a 384 well plate, followed by 4uL of diluted cDNA. All samples were run in technical triplicate. qPCR was then run on a LightCycler 480 (Roche).

### qPCR Analysis

All qPCR primers were validated prior to use in all experiments. All primers were required to generate a single PCR product as determined by melt curve analysis and to have an efficiency of 1.8-2.0. standard curves for all validated primer sets were obtained and used to convert Cp values into relative concentrations. RT-qPCR samples were normalized to housekeeping gene, Rpl13a, then normalized by expression for a non-targeting control normalized by Rpl13a. All qPCR primer sequences are provided in **Table S1**.

### ORCA

The *Fgf5* ORCA probes were designed as previously described^9,51^. We designed ORCA probes to label 38 5kb-regions at 5kb intervals across a 190 kb genomic region encompassing the *Fgf5* regulatory domain and flanking TADs (5kb probe coordinates are provide in **Table S4**).

#### Sample preparation

ORCA imaging experiments were performed following the protocol described^51,52^. Briefly, R1 mESCs expressing CARGO arrays or a non-targeting guide were treated with 2ug/ul doxycycline to induce dCas9-VRP expression for three days. After treatment, cells were fixed in 4% PFA in 1x PBS for 10 min. Cells were washed twice with PBS and spun at 500g for 2 minutes. Cells were resuspended in cold 70% ethanol at a density of 40k cells/uL and stored at -20°C For multiplexed ORCA imaging, each CARGO cell lines were barcoded as previously described^53^. Barcoded cells were combined at desired ratios in a single tube. Cells were plated on a poly-D-lysine-coated 40 mm coverglass and permeabilized with 0.5% Triton-X in 1x PBS for 10 min, followed by 2 washes with 1x PBS. Cells were incubated for 5 minutes in 0.1 M HCl, followed by 3 washes with 1x PBS and 3 washes with 2x SSC. We then treated cells for 35 min in 2x SSC + 50% vol/vol formamide and 0.1% Tween. 3 μg *Fgf5* ORCA probe in 30 μl Hybridization buffer (2x SSC, 50% vol/vol formamide and 0.1% Tween) were added onto cells. Cells were then denatured for 3min at 90°C and incubated overnight at 42°C. After the hybridization, cells were washed twice for 10min in 2x SSC at 42°C, then postfixed in 2% GA, 8% PFA in 1x PBS for 30 min. Cells were then washed in 2x SSC and either imaged directly or stored for up to a week at 4°C.

#### Image acquisition

Samples were imaged on the custom microscopy and microfluidics setup as described^9,51^. Briefly, samples were mounted into a flow chamber. Readout barcodes were visualized sequentially by complementary oligos carrying Cy5 dye followed strand displacement and washes. A Cy3 oligo that label all barcodes were also imaged at each round of imaging as fiducial spots. The fiducial spots were subsequently used for spot calling and registration in the image analysis pipeline.

#### Image processing

Spot calling, localization, and drift correction were performed as previously described^9,51^. Software for processing the raw data is available at https://doi.org/10.5281/zenodo.7698979.

#### ORCA analysis

Processed ORCA data was further filtered, imputed and analyzed in R. Briefly, alleles with fewer than 20 of 38 barcodes imaged were removed. Barcodes that were imaged inconsistently, those that were detected in fewer than 50% of alleles and one that was detected in nearly all alleles, were removed. After calculating pairwise distance matrices for each allele, missing data was imputed as the average distance between neighbors upper and lower and left and right neighbors on the matrix.

Contact frequency between each CARGO array target site and a promoter was calculated from the pairwise contact frequency matrix. The extrapolated contact frequency was calculated as the weighted average of the contact frequencies between each of the two probes nearest the CARGO array and each of the two probes nearest the promoter.

### Generating promoter swap line

Promoter swap lines were generated by sequential replacement of each promoter in which the *Prdm8* promoter was replaced by that of *Fgf5* before the *Prdm8* promoter replaced the endogenous *Fgf5* promoter. mESCs were transfected using lipofectamine 2000 per manufacturers protocol with px458-mCherry carrying Cas9 and a single sgRNA targeting the N-terminus of *Prdm8* and a donor template containing 1kb homology arms, PGK-driven puromycin resistance gene flanked by Frt sites, and the 450bp *Fgf5* promoter. After Puromycin selection, cells were single-cell sorted by FACs and homozygous clones were selected. The selection cassette was removed by transfecting cells using lipofectamine 2000 as above with flippase (addgene #13787). Isolated *Fgf5-Fgf5* clones were then transfected using lipofectamine 2000 as above with px458-mCherry carrying Cas9 and a single sgRNA targeting the N-terminus of *Fgf5* and a donor template containing 1kb homology arms, PGK-driven Neomycin resistance gene flanked by LoxP sites, and the 450bp *Prdm8* promoter. After selection with 200ug/mL G418 (Invivogen) cells were single-cell sorted by FACS and homozygous clones were selected. The selection cassette was then removed by transfected cells using Lipofectamine 2000 with Cre-mCherry. mCherry+ cells were then single-cell sorted and correct genotype was confirmed by PCR and sanger sequencing.

### Generating LTR deletion line

LTR deletion lines were generated using Cas9 and two guides flanking the LTR (LTR-ups 5’-CACCGAGAGATGTGCGTCTGTAATTT-3’, LTR-dns 5’-CACCGAAGCTGTGATTTACCCGAG -3’). mESCs were transfected using lipofectamine 2000 per manufacturers protocol with px458-mCherry carrying Cas9 and one sgRNA and px458-BFP carrying Cas9 and the other guide. Within 48 hours of transfection, mCherry+/BFP+ double positive cells were single cell sorted on a FACS Aria Fusion flow cytometer (BD) in the Stanford Shared FACS facility. The correct genotype was confirmed by PCR and sanger sequencing.

## Supplementary Information

Table S1. Oligonucleotides used in this study

Table S2. Guide sequences and enrichment in CRISPRa Screen

Table S3. CARGO oligonucleotides and guides

Table S4. ORCA probe coordinates

### Supplemental Figures

**Figure S2.**
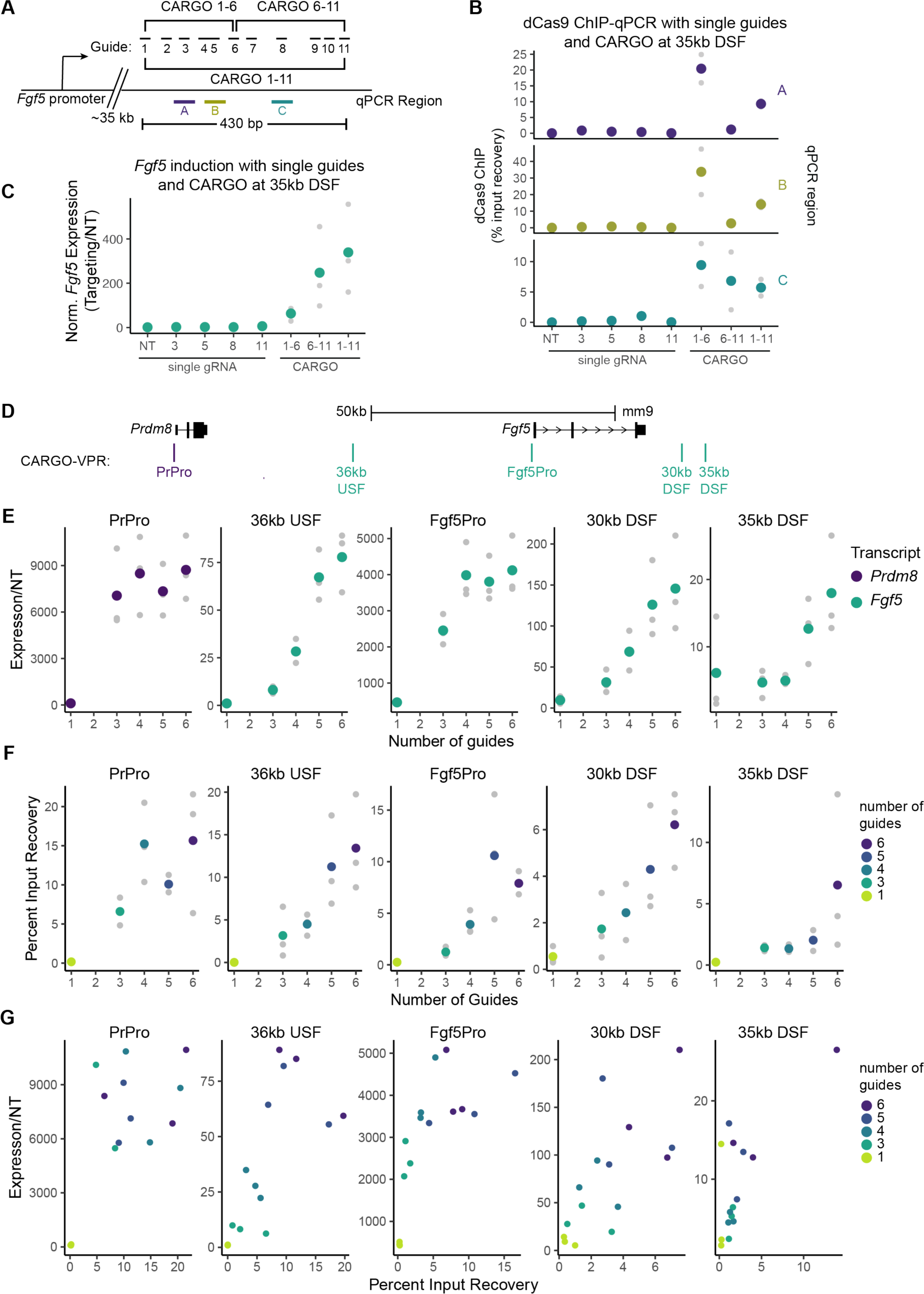
CARGO-VPR enables dCas9 binding at site 35kb downstream from *Fgf5* and induction of *Fgf5* expression. Related to Figure 2. **A.** Positions of eleven individual guides, two 6-mer arrays, and one 11-mer array designed to target a region 35kb DSF. Locations of PCR amplicons used in dCas9 ChIP-qPCR are shown as colored lines labeled A, B, and C. **B**. dCas9 ChIP-qPCR percent input recovery at sites labeled in (B) in cells expressing dCas9-VPR and single guides (NT, 3,5, or 8), 6-mer arrays (1-6 or 6-11), or a 11-mer array (1-11). Colored dots are the mean of two biological replicates and grey dots are the individual replicates. **C.** RT-qPCR of *Fgf5* expression in cells expressing dCas9-VPR and single guides (NT, 3,5, or 8), 6-mer arrays (1-6 or 6-11), or a 11-mer array (1-11). Colored dots are the mean of three biological replicates and grey dots are the individual replicates. **D.** Schematic of dCas9-VPR targeted sites used in guide titration experiments. **E.** Expression measured by rt-qPCR following dCas9-VPR recruitment to designated sites shown on top, using an indicated number of guide RNAs. Expression was normalized to a non-targeting guide control. Color indicates the transcript measured. Colored points are the mean of 3 replicates shown as grey circles. **F.** dCas9-VPR ChIP-qPCR percent input recovery upon dCas9-VPR recruitment using indicated number of guide RNAs. qPCR primers overlap CARGO-VPR target site. Colors indicate number of guide RNAs and are the mean of the 3 replicates shown as grey circles. **G.** Expression/NT from (**E**) plotted against percent input recovery from (**F**) for each of the CARGO-VPR target sites. Colors indicate number of guide RNAs. Each point represents one paired replicate for which rt-qPCR and ChIP qPCR were performed.

**Figure S3.**
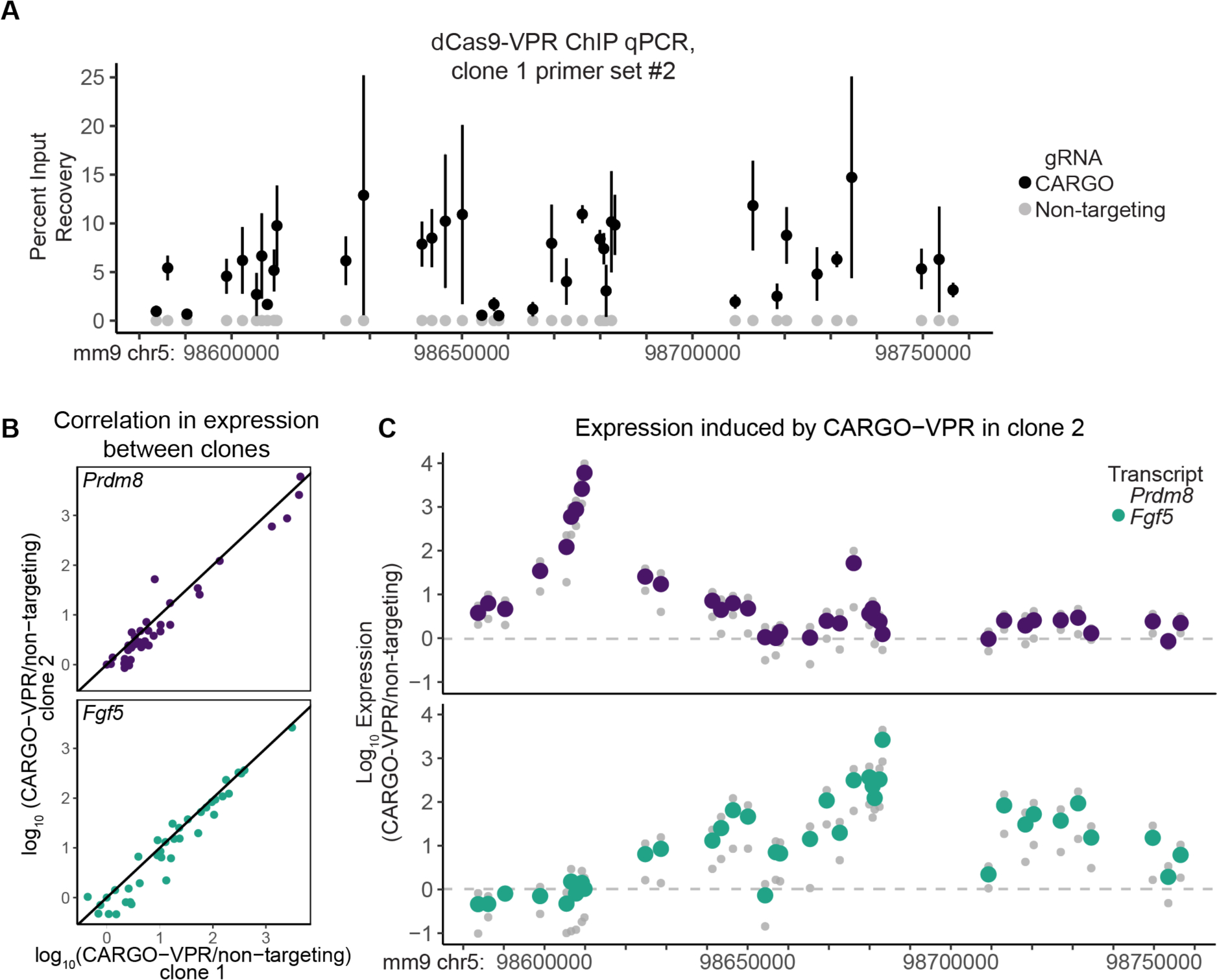
Distance dependent expression decay replicated. Related to Figure 3. **A**. dCas9 ChIP-qPCR in all 38 CARGO populations assayed with a second primer set different than that used in Fig 3C. Black points are the mean of n= 3-4 biological replicates. Error bars are standard deviation. Grey points indicate the same qPCR amplicon assayed from a cell population expressing a non-targeting guide (n=2). **B**. Comparison of *Fgf5* and *Prdm8* expression at each of the CARGO sites in dCas9-VPR clone 2 versus clone 1 (used in Figure 3). Line is y=x. **C**. RT-qPCR of *Prdm8* (upper, purple) and *Fgf5* (lower, green) in cell populations derived from a second, independent dCas9-VPR expressing clonal mESC line to which individual CARGO arrays were introduced. Colored dots are the mean of two replicates. Grey points are means of technical replicates. Expression is plotted as expression normalized to housekeeping gene, *Rpl13a*, over normalized expression from cells expressing a non-targeting guide.

**Figure S4.**
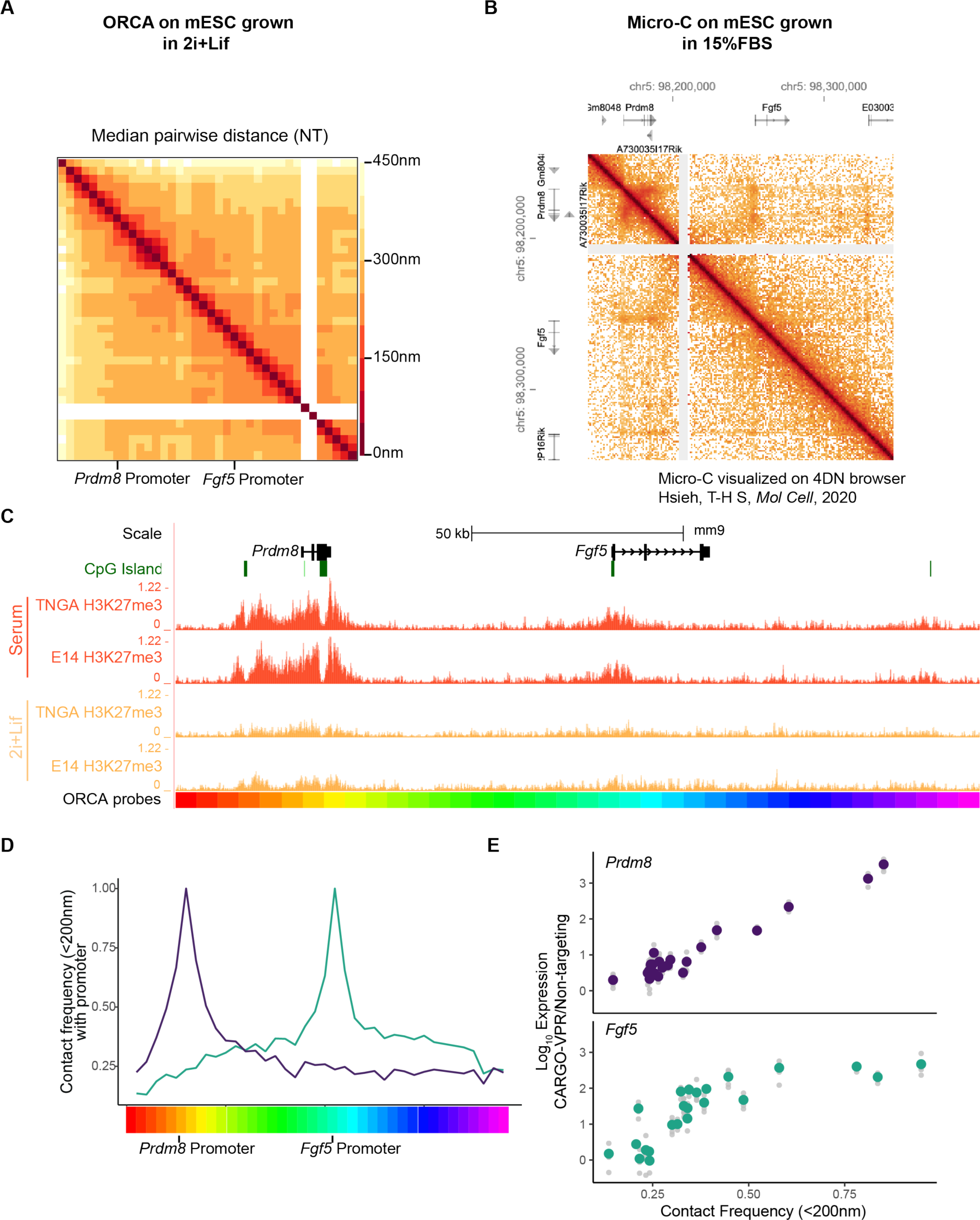
Median distance and contact probability measured at the *Prdm8-Fgf5* locus using ORCA. Related to Figure 4. **A**. Median pairwise distance between 5kb ORCA probes at the *Prdm8-Fgf5* locus in cells grown in 2i+Lif expressing dCas9-VPR and a non-targeting guide (n=2188). White regions at probes 32 and 33 indicate missing data. B. Micro-C from mESCs grown in Serum conditions from Hsieh T-H S, et. al visualized on the 4DN browser^44^ **C**. Genome browser tracks showing annotated CpG islands, and H3K27me3 enrichments in two distinct mESC lines cultured in Serum or 2i+Lif media conditions from Marks et. al^45^. **D**. Pairwise contact probability plots from the probes overlapping the *Prdm8* promoter (purple) or the *Fgf5* promoter (green) and all other probes. Location of individual probes is shown as rainbow-colored blocks and corresponds to placement in (3A). **E.** Log_10_ Normalized Expression of *Prdm8* (upper) and *Fgf5* (lower) induced by CARGO-VPR (y-axis) by contact frequency (x-axis) between the promoter and the CARGO target site, as extrapolated from ORCA measurements. Colored dots are the mean of at least four RT-qPCR replicates. Grey points are individual expression values.

**Figure S5.**
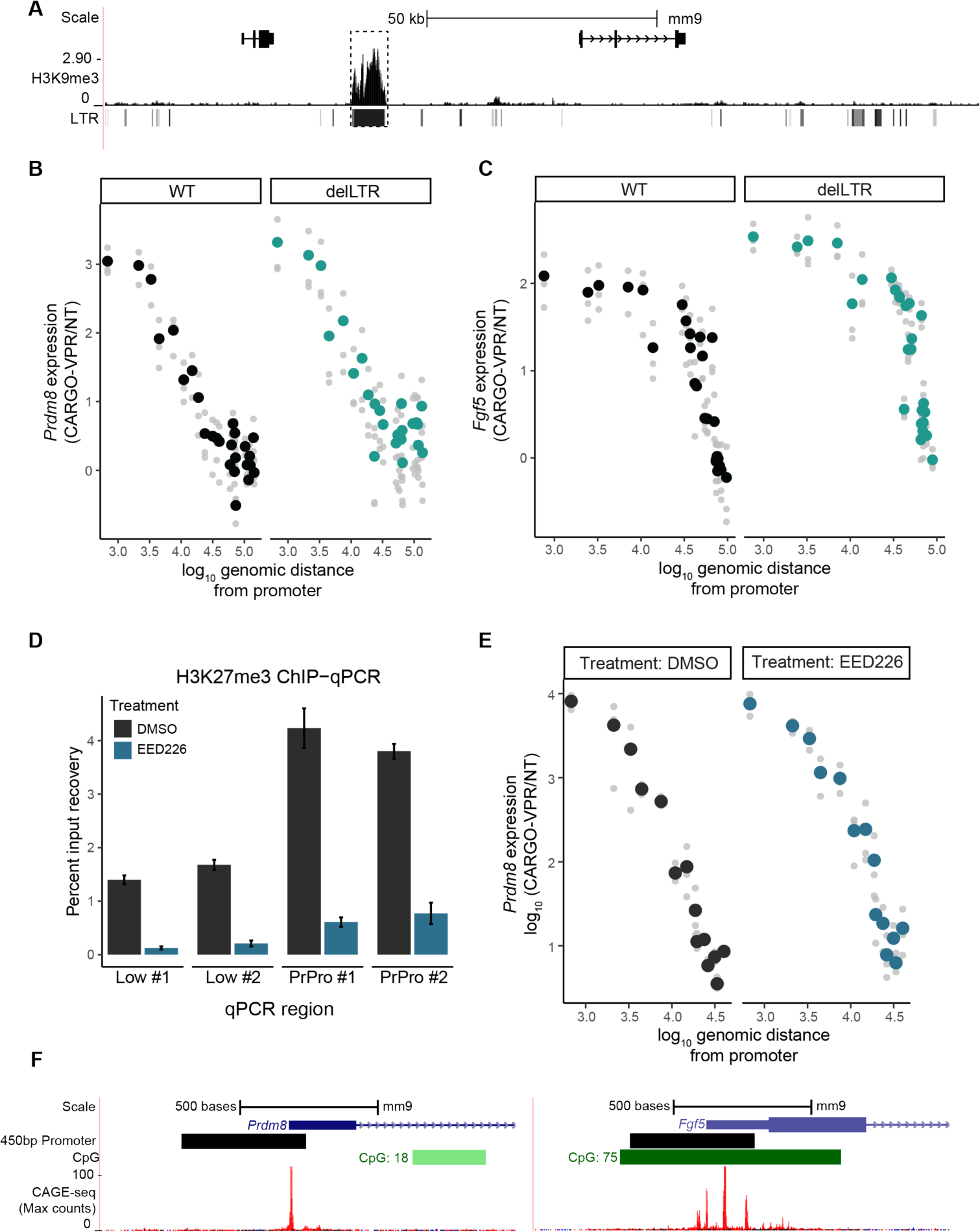
Changing chromatin state of the *Prdm8* promoter does not alter scaling of expression responses. Related to Figure 5. **A.** Browser tracks showing H3K9me3 (ChIP-seq) enrichment^46^ and LTR at the *Prdm8-Fgf5* locus. Deleted LTR element is highlighted with a dashed box. **B.** RT-qPCR for *Prdm8* expression following dCas9-VPR induction in WT or delK9 cells expressing CARGO. Black and teal circles are the mean of three biological replicates and grey points are individual data points **C.** RT-qPCR for *Fgf5* expression following dCas9-VPR induction in WT or delK9 cells expressing CARGO. Black and teal circles are the mean of three biological replicates and grey points are individual data points **D.** H3K27me3 ChIP-qPCR in cells treated with EED226 or vehicle control assayed at downstream sites with low H3K27me3 enrichment (Low #1, Low#2) and at the *Prdm8* promoter (PrPro #1, PrPro #2). Error bars are mean ± standard deviation of n=2 biological replicates. **E**. RT-qPCR for *Prdm8* expression after dCas9-VPR induction in cells expressing CARGO arrays treated with EED226 or vehicle control (DMSO). Black and blue circles are the mean of three biological replicates and grey points are individual data points **F.** Browser tracks showing 450bp promoters, CpG islands, and annotated TSS (FANTOM5 CAGE-seq at the *Prdm8* (left) and *Fgf5* (right) promoters.

## References

1. Banerji, J., Rusconi, S., and Schaffner, W. (1981). Expression of a β-globin gene is enhanced by remote SV40 DNA sequences. Cell 27, 299–308. 10.1016/0092-8674(81)90413-X.

2. Kim, S., and Wysocka, J. (2023). Deciphering the multi-scale, quantitative cis-regulatory code. Mol. Cell 83, 373–392. 10.1016/J.MOLCEL.2022.12.032.

3. Long, H.K., Prescott, S.L., and Wysocka, J. (2016). Ever-changing landscapes: transcriptional enhancers in development and evolution. Cell 167, 1170–1187. 10.1016/j.cell.2016.09.018.

4. Furlong, E.E.M., and Levine, M. (2018). Developmental enhancers and chromosome topology. Science (80-.). 361, 1341–1345. 10.1126/SCIENCE.AAU0320/ASSET/412087A1-DE48-433F-BC76-9CA9B70B9A1B/ASSETS/GRAPHIC/361_1341_F4.JPEG.

5. Reiter, F., Wienerroither, S., and Stark, A. (2017). Combinatorial function of transcription factors and cofactors. Curr. Opin. Genet. Dev. 43, 73–81. 10.1016/J.GDE.2016.12.007.

6. Long, H.K., Osterwalder, M., Welsh, I.C., Hansen, K., Davies, J.O.J., Liu, Y.E., Koska, M., Adams, A.T., Aho, R., Arora, N., et al. (2020). Loss of extreme long-range enhancers in human neural crest drives a craniofacial disorder. Cell Stem Cell 27, 765–783.e14. 10.1016/j.stem.2020.09.001.

7. Bahr, C., Von Paleske, L., Uslu, V. V., Remeseiro, S., Takayama, N., Ng, S.W., Murison, A., Langenfeld, K., Petretich, M., Scognamiglio, R., et al. (2018). A Myc enhancer cluster regulates normal and leukaemic haematopoietic stem cell hierarchies. Nature 553, 515–520. 10.1038/nature25193.

8. Lettice, L.A., Heaney, S.J.H., Purdie, L.A., Li, L., de Beer, P., Oostra, B.A., Goode, D., Elgar, G., Hill, R.E., and de Graaff, E. (2003). A long-range Shh enhancer regulates expression in the developing limb and fin and is associated with preaxial polydactyly. Hum. Mol. Genet. 12, 1725–1735. 10.1093/hmg/ddg180.

9. Mateo, L.J., Murphy, S.E., Hafner, A., Cinquini, I.S., Walker, C.A., and Boettiger, A.N. (2019). Visualizing DNA folding and RNA in embryos at single-cell resolution. Nat. 2019 5687750 568, 49–54. 10.1038/S41586-019-1035-4.

10. Alexander, J.M., Guan, J., Li, B., Maliskova, L., Song, M., Shen, Y., Huang, B., Lomvardas, S., and Weiner, O.D. (2019). Live-cell imaging reveals enhancer-dependent Sox2 transcription in the absence of enhancer proximity. Elife 8. 10.7554/ELIFE.41769.

11. Chen, H., Levo, M., Barinov, L., Fujioka, M., Jaynes, J.B., and Gregor, T. (2018). Dynamic interplay between enhancer–promoter topology and gene activity. Nat. Genet. 2018 509 50, 1296–1303. 10.1038/s41588-018-0175-z.

12. Li, J., Dong, A., Saydaminova, K., Chang, H., Wang, G., Ochiai, H., Yamamoto, T., and Pertsinidis, A. (2019). Single-Molecule Nanoscopy Elucidates RNA Polymerase II Transcription at Single Genes in Live Cells. Cell. 10.1016/j.cell.2019.05.029.

13. Li, J., Hsu, A., Hua, Y., Wang, G., Cheng, L., Ochiai, H., Yamamoto, T., and Pertsinidis, A. (2020). Single-gene imaging links genome topology, promoter–enhancer communication and transcription control. Nat. Struct. Mol. Biol. 27, 1032–1040. 10.1038/s41594-020-0493-6.

14. Benabdallah, N.S., Williamson, I., Illingworth, R.S., Kane, L., Boyle, S., Sengupta, D., Grimes, G.R., Therizols, P., and Bickmore, W.A. (2019). Decreased Enhancer-Promoter Proximity Accompanying Enhancer Activation. Mol. Cell 76, 473–484.e7. 10.1016/J.MOLCEL.2019.07.038.

15. Zuin, J., Roth, G., Zhan, Y., Cramard, J., Redolfi, J., Piskadlo, E., Mach, P., Kryzhanovska, M., Tihanyi, G., Kohler, H., et al. (2022). Nonlinear control of transcription through enhancer–promoter interactions. Nat. 2022 6047906 604, 571–577. 10.1038/S41586-022-04570-Y.

16. Xiao, J., Hafner, A., and Boettiger, A.N. (2021). How subtle changes in 3D structure can create large changes in transcription. Elife 10. 10.7554/ELIFE.64320.

17. Nora, E.P., Lajoie, B.R., Schulz, E.G., Giorgetti, L., Okamoto, I., Servant, N., Piolot, T., van Berkum, N.L., Meisig, J., Sedat, J., et al. (2012). Spatial partitioning of the regulatory landscape of the X-inactivation centre. Nature 485, 381–385. 10.1038/nature11049.

18. Dixon, J.R., Selvaraj, S., Yue, F., Kim, A., Li, Y., Shen, Y., Hu, M., Liu, J.S., and Ren, B. (2012). Topological domains in mammalian genomes identified by analysis of chromatin interactions. Nature 485, 376–380. 10.1038/nature11082.

19. Symmons, O., Uslu, V.V., Tsujimura, T., Ruf, S., Nassari, S., Schwarzer, W., Ettwiller, L., and Spitz, F. (2014). Functional and topological characteristics of mammalian regulatory domains. Genome Res. 24, 390–400. 10.1101/GR.163519.113.

20. Chakraborty, S., Kopitchinski, N., Zuo, Z., Eraso, A., Awasthi, P., Chari, R., Mitra, A., Tobias, I.C., Moorthy, S.D., Dale, R.K., et al. (2023). Enhancer–promoter interactions can bypass CTCF-mediated boundaries and contribute to phenotypic robustness. Nat. Genet. 2023 552 55, 280–290. 10.1038/s41588-022-01295-6.

21. Hung, T.-C., Kingsley, D.M., and Boettiger, A. (2023). Boundary stacking interactions enable cross-TAD enhancer-promoter communication during limb development. bioRxiv, 2023.02.06.527380.

22. Pombo, A., and Nicodemi, M. (2014). Physical mechanisms behind the large scale features of chromatin organization. Transcription 5. 10.4161/TRNS.28447.

23. Fulco, C.P., Nasser, J., Jones, T.R., Munson, G., Bergman, D.T., Subramanian, V., Grossman, S.R., Anyoha, R., Doughty, B.R., Patwardhan, T.A., et al. (2019). Activity-by-contact model of enhancer–promoter regulation from thousands of CRISPR perturbations. Nat. Genet. 2019 5112 51, 1664–1669. 10.1038/s41588-019-0538-0.

24. Fuentes, D.R., Swigut, T., and Wysocka, J. (2018). Systematic perturbation of retroviral LTRs reveals widespread long-range effects on human gene regulation. Elife 7. 10.7554/eLife.35989.

25. Yokoshi, M., Segawa, K., and Fukaya, T. (2020). Visualizing the Role of Boundary Elements in Enhancer-Promoter Communication. Mol. Cell 0. 10.1016/j.molcel.2020.02.007.

26. Rinzema, N.J., Sofiadis, K., Tjalsma, S.J.D., Verstegen, M.J.A.M., Oz, Y., Valdes-Quezada, C., Felder, A.K., Filipovska, T., van der Elst, S., de Andrade dos Ramos, Z., et al. (2022). Building regulatory landscapes reveals that an enhancer can recruit cohesin to create contact domains, engage CTCF sites and activate distant genes. Nat. Struct. Mol. Biol. 2022 296 29, 563–574. 10.1038/S41594-022-00787-7.

27. Brückner, D.B., Chen, H., Barinov, L., Zoller, B., and Gregor, T. (2023). Stochastic motion and transcriptional dynamics of pairs of distal DNA loci on a compacted chromosome. Science (80-.). 380, 1357–1362. 10.1126/SCIENCE.ADF5568/SUPPL_FILE/SCIENCE.ADF5568_MDAR_REPRODUCIBILITY_CHECKLIST.PDF.

28. Haberle, V., Arnold, C.D., Pagani, M., Rath, M., Schernhuber, K., and Stark, A. (2019). Transcriptional cofactors display specificity for distinct types of core promoters. Nature 570, 122–126. 10.1038/s41586-019-1210-7.

29. Kane, L., Williamson, I., Flyamer, I.M., Kumar, Y., Hill, R.E., Lettice, L.A., and Bickmore, W.A. (2022). Cohesin is required for long-range enhancer action at the Shh locus. Nat. Struct. Mol. Biol. 29, 891–897. 10.1038/S41594-022-00821-8.

30. Hilton, I.B., D’Ippolito, A.M., Vockley, C.M., Thakore, P.I., Crawford, G.E., Reddy, T.E., and Gersbach, C.A. (2015). Epigenome editing by a CRISPR-Cas9-based acetyltransferase activates genes from promoters and enhancers. Nat. Biotechnol. 33, 510–517. 10.1038/NBT.3199.

31. Kuscu, C., Mammadov, R., Czikora, A., Unlu, H., Tufan, T., Fischer, N.L., Arslan, S., Bekiranov, S., Kanemaki, M., and Adli, M. (2019). Temporal and Spatial Epigenome Editing Allows Precise Gene Regulation in Mammalian Cells. J. Mol. Biol. 431, 111–121. 10.1016/j.jmb.2018.08.001.

32. Li, K., Liu, Y., Cao, H., Zhang, Y., Gu, Z., Liu, X., Yu, A., Kaphle, P., Dickerson, K.E., Ni, M., et al. (2020). Interrogation of enhancer function by enhancer-targeting CRISPR epigenetic editing. Nat. Commun. 11. 10.1038/S41467-020-14362-5.

33. Boulos, J., Ehrlich, I., Avidan, N., Shkedi, O., Haimovich-Caspi, L., Kaplan, N., and Kehat, I. (2022). Recruitment of transcriptional effectors by Cas9 creates cis regulatory elements and demonstrates distance-dependent transcriptional regulation. bioRxiv, 2022.02.03.478957. 10.1101/2022.02.03.478957.

34. Alerasool, N., Leng, H., Lin, Z.Y., Gingras, A.C., and Taipale, M. (2022). Identification and functional characterization of transcriptional activators in human cells. Mol. Cell 82, 677–695.e7. 10.1016/J.MOLCEL.2021.12.008.

35. Gu, B., Swigut, T., Spencley, A., Bauer, M.R., Chung, M., Meyer, T., and Wysocka, J. (2018). Transcription-coupled changes in nuclear mobility of mammalian cis-regulatory elements. Science (80-.). *359*, 1050–1055. 10.1126/science.aao3136.

36. Yan, J., Chen, S.A.A., Local, A., Liu, T., Qiu, Y., Dorighi, K.M., Preissl, S., Rivera, C.M., Wang, C., Ye, Z., et al. (2018). Histone H3 lysine 4 monomethylation modulates long-range chromatin interactions at enhancers. Cell Res. 2018 282 28, 204–220. 10.1038/cr.2018.1.

37. Buecker, C., Srinivasan, R., Wu, Z., Calo, E., Acampora, D., Faial, T., Simeone, A., Tan, M., Swigut, T., and Wysocka, J. (2014). Reorganization of Enhancer Patterns in Transition from Naive to Primed Pluripotency. Cell Stem Cell 14, 838– 853. 10.1016/J.STEM.2014.04.003..

38. Thomas, H.F., Kotova, E., Jayaram, S., Pilz, A., Romeike, M., Lackner, A., Penz, T., Bock, C., Leeb, M., Halbritter, F., et al. (2021). Temporal dissection of an enhancer cluster reveals distinct temporal and functional contributions of individual elements. Mol. Cell 81, 969–982.e13. 10.1016/J.MOLCEL.2020.12.047..

39. Konermann, S., Brigham, M.D., Trevino, A.E., Joung, J., Abudayyeh, O.O., Barcena, C., Hsu, P.D., Habib, N., Gootenberg, J.S., Nishimasu, H., et al. (2014). Genome-scale transcriptional activation by an engineered CRISPR-Cas9 complex. Nature 517. 10.1038/nature14136.

40. Pulecio, J., Tayyebi, Z., Liu, D., Wong, W., Luo, R., Damodaran, J.R., Kaplan, S., Cho, H., Yan, J., Murphy, D., et al. (2023). Discovery of Competent Chromatin Regions in Human Embryonic Stem Cells. bioRxiv.

41. Horlbeck, M.A., Witkowsky, L.B., Guglielmi, B., Replogle, J.M., Gilbert, L.A., Villalta, J.E., Torigoe, S.E., Tjian, R., and Weissman, J.S. (2016). Nucleosomes impede Cas9 access to DNA in vivo and in vitro. Elife 5. 10.7554/ELIFE.12677..

42. Isaac, R.S., Jiang, F., Doudna, J.A., Lim, W.A., Narlikar, G.J., and Almeida, R. Nucleosome breathing and remodeling constrain CRISPR-Cas9 function. 10.7554/eLife.13450.001.

43. Dorighi, K.M., Swigut, T., Henriques, T., Bhanu, N. V., Scruggs, B.S., Nady, N., Still, C.D., Garcia, B.A., Adelman, K., and Wysocka, J. (2017). Mll3 and Mll4 Facilitate Enhancer RNA Synthesis and Transcription from Promoters Independently of H3K4 Monomethylation. Mol. Cell. 10.1016/j.molcel.2017.04.018.

44. Hsieh, T.H.S., Cattoglio, C., Slobodyanyuk, E., Hansen, A.S., Rando, O.J., Tjian, R., and Darzacq, X. (2020). Resolving the 3D Landscape of Transcription-Linked Mammalian Chromatin Folding. Mol. Cell 78, 539–553.e8. 10.1016/J.MOLCEL.2020.03.002.

45. Marks, H., Kalkan, T., Menafra, R., Denissov, S., Jones, K., Hofemeister, H., Nichols, J., Kranz, A., Francis Stewart, A., Smith, A., et al. (2012). The Transcriptional and Epigenomic Foundations of Ground State Pluripotency. Cell 149, 590. 10.1016/J.CELL.2012.03.026.

46. Spencley, A.L., Bar, S., Swigut, T., Flynn, R.A., Lee, C.H., Chen, L.F., Bassik, M.C., and Wysocka, J. (2023). Co-transcriptional genome surveillance by HUSH is coupled to termination machinery. Mol. Cell 83, 1623–1639.e8. 10.1016/j.molcel.2023.04.014.

47. Symmons, O., Pan, L., Remeseiro, S., Aktas, T., Klein, F., and Huber, W. (2016). The Shh Topological Domain Facilitates the Action of Remote Enhancers by Reducing the Effects of Genomic Distances. Dev. Cell 39, 529–543. 10.1016/j.devcel.2016.10.015.

48. Batut, P.J., Bing, X.Y., Sisco, Z., Raimundo, J., Levo, M., and Levine, M.S. (2022). Genome organization controls transcriptional dynamics during development. Science (80-.). *375*. 10.1126/SCIENCE.ABI7178.

49. Thomas, H.F., Kotova, E., Jayaram, S., Pilz, A., Romeike, M., Lackner, A., Penz, T., Bock, C., Leeb, M., Halbritter, F., et al. (2021). Temporal dissection of an enhancer cluster reveals distinct temporal and functional contributions of individual elements. Mol. Cell 81, 969–982.e13. 10.1016/J.MOLCEL.2020.12.047.

50. Buecker, C., Srinivasan, R., Wu, Z., Calo, E., Acampora, D., Faial, T., Simeone, A., Tan, M., Swigut, T., and Wysocka, J. (2014). Reorganization of enhancer patterns in transition from naive to primed pluripotency. Cell Stem Cell 14, 838– 853. 10.1016/j.stem.2014.04.003.

51. Mateo, L.J., Sinnott-Armstrong, N., and Boettiger, A.N. (2021). Tracing DNA paths and RNA profiles in cultured cells and tissues with ORCA. Nat. Protoc. 2021 163 16, 1647–1713. 10.1038/s41596-020-00478-x.

52. Bintu, B., Mateo, L.J., Su, J.H., Sinnott-Armstrong, N.A., Parker, M., Kinrot, S., Yamaya, K., Boettiger, A.N., and Zhuang, X. (2018). Super-resolution chromatin tracing reveals domains and cooperative interactions in single cells. Science (80-.). 362. 10.1126/SCIENCE.AAU1783.

53. Hafner, A., Park, M., Berger, S.E., Murphy, S.E., Nora, E.P., and Boettiger, A.N. (2023). Loop stacking organizes genome folding from TADs to chromosomes. Mol. Cell 83, 1377–1392.e6. 10.1016/J.MOLCEL.2023.04.008.

